# LIN-15B promotes enrichment of H3K9me2 on the promoters of a subset of germline genes that are repressed in somatic cells in *C. elegans*

**DOI:** 10.1101/497438

**Authors:** Andreas Rechtsteiner, Meghan E. Costello, Thea A. Egelhofer, Jacob M. Garrigues, Susan Strome, Lisa N. Petrella

## Abstract

Repression of germline-promoting genes in somatic cells is critical for somatic development and function. To study how germline genes are repressed in somatic tissues, we analyzed key histone modifications in three *Caenorhabditis elegans* synMuv B mutants, *lin-15B, lin-35,* and *lin-37,* all of which display ectopic expression of germline genes in the soma. LIN-35 and LIN-37 are members of the conserved DREAM complex. LIN-15B has been proposed to work with the DREAM complex but has not been shown biochemically to be a complex member. We found that in wild-type worms synMuv B target genes and germline genes are enriched for the repressive histone modification dimethylation of histone H3 on lysine 9 (H3K9me2) at their promoters. Genes with H3K9me2 promoter localization are distributed across the autosomes and not biased toward autosomal arms like broad H3K9me2 domains. Both synMuv B targets and germline genes display dramatic reduction of H3K9me2 promoter localization in *lin-15B* mutants, but much weaker reduction in *lin-35* and *lin-37* mutants. This is the first major difference reported between *lin-15B* and DREAM complex mutants, which likely represents a difference in molecular function for these synMuv B proteins. In support of the pivotal role of H3K9me2 in regulation of germline genes through LIN-15B, global loss of H3K9me2 but not H3K9me3 results in phenotypes similar to synMuv B mutants, high temperature larval arrest and ectopic expression of germline genes in the soma. We propose that LIN-15B-driven enrichment of H3K9me2 on promoters of germline genes contributes to repression of those genes in somatic tissues.

## INTRODUCTION

Repression in somatic cells of genes that promote germline development and function is a vital cell fate regulatory mechanism, which when disrupted leads to developmental problems and is a hallmark of aggressive cancer (Janic *et al.* 2010; Petrella *et al.* 2011; Whitehurst 2014; Al-Amin *et al.* 2016). Repression of germline genes in the soma poses a unique challenge for cells. First, like other genes expressed in specific tissues, germline genes can be found clustered along chromosomes; however, within a given cluster genes with ubiquitous, germline-specific, germline-enriched, and non-germline expression are interspersed (Spellman and Rubin 2002; Roy *et al.* 2002; Reinke and Cutter 2009).

Therefore, somatic cells require a mechanism to repress germline-specific genes without disrupting expression of important flanking genes. Second, because embryos start life as the fusion of two germline cells, an egg and a sperm, they inherit an epigenetic state that has been driving germline gene expression (Furuhashi *et al.* 2010; Rechtsteiner *et al.* 2010; Zenk *et al.* 2017; Tabuchi *et al.* 2018; Kreher *et al.* 2018). This chromatin state must be reset during development to turn off germline gene expression in differentiating somatic cells (Morgan *et al.* 2005; Fraser and Lin 2016). There has been no investigation to date of the unique patterns of chromatin modifications or regulatory protein binding that lead to repression of germline-specific genes across somatic tissues.

synMuv (for synthetic Multivulva) B proteins are a diverse class of transcriptional repressors that are involved in a number of different fate decisions in *C. elegans* (Unhavaithaya *et al.* 2002; Wang *et al.* 2005; Fay and Yochem 2007). A subset of synMuv B genes show a distinct set of mutant phenotypes, which include ectopic expression of germline genes in somatic cells and larval arrest at high temperature (called HTA for high temperature arrest) (Wang *et al.* 2005; Petrella *et al.* 2011; Wu *et al.* 2012). Of this subset, a large proportion encode proteins that exist in two proposed complexes: the HP1-containing heterochromatin complex (HPL-2, LIN-13, LIN-61 and the DREAM complex (EFL-1, DPL-2, LIN-35, LIN-9, LIN-37, LIN-52, LIN-53, LIN-54) (Coustham *et al.* 2006; Harrison *et al.* 2006; Wu *et al.* 2012). Several additional synMuv B mutants, including *lin-15B* and *met-2,* also display ectopic germline gene expression in the soma, but have not been shown biochemically to encode members of the HP1 or DREAM complex (Wu *et al.* 2012; Petrella *et al.* 2011). *lin-15B* mutants, like mutants in genes encoding DREAM complex members, also display an HTA phenotype, show changes in regulation of somatic RNAi, and cause transgene silencing in the soma (Wang *et al.* 2005; Petrella *et al.* 2011; Wu *et al.* 2012).

While mutations in genes encoding the heterochromatin complex, the DREAM complex, LIN-15B, and MET-2 all lead to ectopic expression of germline genes in the soma, the precise way these different complexes/proteins function in parallel or together to repress germline genes in the somatic tissues of wild-type animals is not understood.

Several lines of evidence point to synMuv B complexes repressing gene expression by altering chromatin. First, synMuv B mutant phenotypes, including HTA and ectopic germline gene expression, are strongly suppressed by loss of chromatin factors (Unhavaithaya *et al.* 2002; Wang *et al.* 2005; Cui *et al.* 2006; Petrella *et al.* 2011; Wu *et al.* 2012). Second, the DREAM complex has been shown to promote enrichment of the H2A histone variant HTZ-1 in the body of a subset of genes that the DREAM complex represses in L3 larvae (Latorre *et al.* 2015). Finally, HPL-2 is a homolog of heterochromatin protein 1 (HP1) (Couteau *et al.* 2002). HPL-2, in a complex with LIN-13 and LIN-61, localizes to areas of histone H3 methylated at lysine 9 (H3K9me) and helps create repressive heterochromatin (Wu *et al.* 2012; Garrigues *et al.* 2015). Together these data indicate that changes to chromatin may underlie the ectopic expression of germline genes in synMuv B mutants.

One of the best studied aspects of chromatin regulation is covalent modifications on histone tails. Specific histone modifications are often associated with repressive or active chromatin compartments and can be a read-out of the expression state of a gene. Histone H3 lysine 4 methylation (H3K4me) and H3 lysine 36 methylation (H3K36me) are generally associated with areas of previous or active gene expression (Ho *et al.* 2014; Evans *et al.* 2016). In contrast, histone H3 lysine 9 methylation (H3K9me) and histone H3 lysine 27 methylation (H3K27me) are associated with areas of low/no expression of coding genes and repression of repetitive elements (Ahringer and Gasser 2018). Of particular interest for the regulation of germline gene expression in somatic cells is histone H3K9 methylation. In *C. elegans,* mono- and dimethylation of H3K9 (H3K9me1 and H3K9me2, respectively), are primarily catalyzed by MET-2. *met-2* mutants lose 80-90% of H3K9me1 and H3K9me2 in embryos (Towbin *et al.* 2012). *met-2* is a synMuv B gene and mutants have been previously shown to ectopically express germline genes in somatic cells (Wu *et al.* 2012).

Trimethlyation of H3K9 (H3K9me3) is catalyzed by a separate histone methyltransferase, SET-25 (Towbin *et al.* 2012). *set-25* is not a synMuv B gene and its potential role in regulating germline gene expression in the soma has not been tested. Several studies have analyzed the roles in *C. elegans* of H3K9me2 and H3K9me3 in regulating the interaction of heterochromatin with the nuclear periphery and repression of repetitive elements (Meister *et al.* 2010; Towbin *et al.* 2012; Guo *et al.* 2015; Zeller *et al.* 2016). Both of these functions primarily rely on the high enrichment of H3K9 methylation on the heterochromatic arms of the autosomes (Ikegami *et al.* 2010; Liu *et al.* 2011; Garrigues *et al.* 2015; Evans *et al.* 2016). However, little work has been done to look at how H3K9 methylation localizes to or regulates protein-coding genes in the euchromatic central regions of autosomes, especially germline genes. To fill this gap, we sought to determine the changes in the levels and distributions of active and repressive histone modifications in the soma of synMuv B mutants and whether such changes may underlie ectopic expression of germline genes.

In this study we used chromatin immunoprecipitation with genome-wide high-throughput sequencing (ChIP-seq) to investigate histone modifications in wild type and three synMuv B mutants, *lin-15B, lin-35,* and *lin-37.* We found that in wild-type L1 larvae, which are composed of 550 somatic cells and 2 germ cells and are therefore primarily somatic, H3K9me2 is enriched on the promoters of a subset of genes that display germline-specific expression. The genes that have H3K9me2 in their promoters in wild type are generally up-regulated in synMuv B mutants, suggesting that H3K9me2 plays a role in their repression. In support of this, the localization of H3K9me2 to gene promoters is largely lost in *lin-15B* mutants and is diminished but not lost in *lin-35* and *lin-37* mutants. Loss of H3K9me2 on promoters in mutants is associated with an increase in H3K4me3 on promoters and H3K36me3 in gene bodies, modifications associated with gene expression, suggesting that these genes go from a repressed to expressed state. Finally, we found that global loss of H3K9me2 but not H3K9me3 results in both the HTA and the ectopic germline gene expression phenotypes seen in *lin-15B* mutants. We propose that LIN-15B and DREAM repress a subset of germline genes in somatic tissues by promoting enrichment of H3K9me2 on those genes’ promoters.

## MATERIALS AND METHODS

### C. elegans strains and culture conditions

*C. elegans* were cultured using standard conditions (Brenner 1974) at 20°C unless otherwise noted. N2 (Bristol) was used as wild type. Mutant strains were as follows:

MT10430 *lin-35(n745) I*

SS1110 *hpl-2(tm1489) III*

MT5470 *lin-37(n758) III*

MT13293 *met-2(n4256) III*

MT17463 *set-25(n5032) III*

LNP0036 *met-2(n4256) set-25(n5032) III*

MT2495 *lin-15B(n744) X*

### ChlP-seq from L1s

Worms were grown from synchronized L1s in standard S-basal medium with shaking at 230 rpm and fed HB101 bacteria until gravid. Embryos were harvested using standard bleaching methods, and L1s were synchronized in S-basal media with shaking for 14-18 hours in the absence of food. For 26°C samples, worms were grown to the L4 stage at 20°C, when they were up-shifted to 26°C until gravid and L1s were harvested as described above. Extracts were made as described in (Kolasinska-Zwierz *et al.* 2009) with the following modifications. Cross-linked chromatin was sonicated using a Diagenode Bioruptor at high setting for 30 pulses, each lasting 30 sec followed by a 1 min pause. ChIP was performed as described by (Kolasinska-Zwierz *et al.* 2009) with the modification of using 0.5 mg of protein and 1 μg antibody or by using an IP-Star Compact Automated System (Diagenode) as described in (Tabuchi *et al.* 2018). Sequencing libraries were prepared in two ways. Some libraries were prepared with the NEBNext Ultra DNA library Prep Kit (NEB) following the manufacturers’ instructions. 1 ng of starting DNA was used, adapters were diluted 1:40, and AMPure beads were used for size selection before amplification to enrich for fragments corresponding to a 200 bp insert size. The other libraries were prepared using Illumina Truseq adapters and primers with the following custom mixtures. ChIP or input DNA fragments were end repaired with the following: 5 μl T4 DNA ligase buffer with 10 mM ATP, 2 μl dNTP mix, 1.2 μl T4 DNA polymerase (3U/μl), 0.8 μl 1:5 Klenow DNA polymerase (diluted with 1X T4 DNA ligase buffer for final Klenow concentration 1U/μl), 1 μl T4 PNK (10U/μl). This 50 μl reaction was incubated at 20°C for 30 minutes and purified with a QIAquick PCR spin column (elution volume 36 μl). ‘A’ bases were then added to the 3’ end of the DNA fragments with the following: 5 μl Klenow buffer/NEB 2, 10 μl dATP (1 mM), 1 μl Klenow 3’ to 5’ exo-(5U/μl).

This mixture was incubated at 37°C for 30 min, and the DNA was purified with a QIAquick MinElute column (11 μl of DNA was eluted into a siliconized tube). Illumina TruSeq adapters were ligated to DNA fragments with the following: 15 μl 2x Rapid Ligation buffer, 1 μl adapters (diluted 1:40), 1.5 μl Quick T4 DNA Ligase. This 30 μl reaction was incubated at 23°C for 30 min. The mixture was then cleaned up 2X with AMPure beads (using 95% volume beads) and DNA was eluted in 22 μl. The Adapter-Modified DNA fragments were amplified by PCR with the following mixture: 6 μl 5X Phusion Buffer HF, 2 μl Primer cocktail (from TrueSeq kit), 0.5 μl 25 mM dNTP mix, 0.5 Phusion polymerase (2U/μl) using the following PCR program: 98°C 30 min, 98°C 10 min and 60°C 30 min and 72°C 30 min repeated 16 cycles, 72°C 5 min. The amplified DNA was concentrated and loaded onto a 2% agarose gel, and DNA between 300-400 bp was recovered from the gel. The multiplexed libraries were sequenced on an Illumina HiSeq4000 or HiSeq2500 at the Vincent J. Coates Genomics Sequencing Laboratory at University of California, Berkeley.

### ChIP-chip from embryos

Late-stage embryos were obtained and chromatin extracts prepared as described in (Latorre *et al.* 2015). Chromatin immunoprecipitation and subsequent LM-PCR, microarray hybridization, and scanning were performed as in (Garrigues *et al.* 2015).

### Antibodies used for ChIP

Mouse monoclonal antibodies for H3K9me2 (Wako MABI0307, #3023069), H3K36me3 (Wako MABI0333, #300–95289), H3K27me3 (Wako MABI0323, #309–95259), and H3K4me3 (Wako MABI0304, #305–34819) as used in (Liu *et al.* 2011; Egelhofer *et al.* 2011). Rabbit polyclonal LIN-15B antibody (SDQ2330, Novus #38610002) was used at a concentration of 2.5 μg per mg of chromatin extract.

### Analysis of ChlP-seq data

Raw sequence reads from the Illumina HiSeq (50 bp single-end reads) were mapped to the *C. elegans* genome (Wormbase version WS220) using Bowtie with default settings (Langmead *et al.* 2009). MACS2 (Zhang *et al.* 2008) was used to call peaks and create bedgraph files for sequenced and mapped H3K4me3 ChIP samples and corresponding Input DNA samples with the following parameters: callpeak -t H3K4me3.mapped.reads.sampleX -c Input.mapped.reads.sampleX -g ce --bdg -- keep-dup=auto --qvalue=0.01 --nomodel --extsize=250 —call-summits

MACS2 was used to call peaks and create bedgraph files for sequenced and mapped H3K9me2 ChIP samples and corresponding Input DNA samples with slightly different parameters to account for the broader domains of H3K9me2: callpeak -t H3K9me2.mapped.reads.sampleX -c Input.mapped.reads.sampleX -g ce --bdg -- keep-dup=auto --broad --broad-cutoff=0.01 --nomodel --extsize=250

A peak was considered to be associated with a gene’s promoter if it overlapped at least 100bp with the region 750bp upstream from the gene’s TSS to 250bp downstream from the TSS. A peak was considered to be associated with the body of a gene if it overlapped at least 250bp with the region from 250bp downstream from the TSS to the TES. A gene’s promoter or gene body was considered bound by H3K4me3 or H3K9me2 in one of the conditions if for all replicates of that condition a peak was associated with the gene’s promoter or body, respectively. The distribution of genes with peaks in promoters or gene bodies along an autosome are shown in Fig. 3A in 200kb windows.

Bedgraph files for genome browser displays were scaled to 5 million total reads for all H3K4me3 ChIP samples, 10 million reads for all H3K36me3 samples, 15 million reads for all H3K9me2 samples, and 20 million reads for all H3K27me3 samples. The different scaling factors roughly correspond to the different genome-wide coverages of the different ChIP factors, e.g. H3K4me3 being found mostly on promoters of expressed genes, H3K36me3 mostly on gene bodies of expressed genes, and H3K9me2 mostly on chromosomal arms. Further data analysis below was based on these scaled read coverages. Scaled bedgraph files were converted to bigwig using the bedGraphToBigWig UCSC Genome Browser tool (Kent *et al.* 2010) and displayed on the UCSC Genome Browser.

### Analysis of LIN-15B ChIP-chip data

NimbleGen 2.1M probe tiling arrays (DESIGN_ID = 8258), with 50 bp probes, designed against WS170 (ce4) were used. Two independent ChIPs were performed. Amplified samples were labeled and hybridized by the Roche NimbleGen Service Laboratory. ChIP samples were labeled with Cy5 and their input reference with Cy3. For each probe, the intensity from the sample channel was divided by the reference channel and log2 transformed. The enrichment scores for each replicate were calculated by standardizing the log ratios to mean zero and standard deviation one (z-score) and the average z-score across replicates was calculated and displayed in the UCSC Genome Browser (Fig S3). Peak calling was performed with the MA2C algorithm (Song *et al.* 2007) using Nimblegen array design files 080922_modEncode_CE_chip_HX1.pos and 080922_modEncode_CE_chip_HX1.ndf and MA2C parameters METHOD = Robust, C = 2, pvalue = 1e-5, BANDWIDTH = 300, MIN_PROBES = 5, MAX_GAP = 250. The resulting peak calls were associated with gene promoters and bodies as described in the previous section.

### Correlation heatmap of samples

The scaled bedgraph files were used to calculate for each sample the average read coverage in 1kb windows across all autosomes and the X chromosome. The resulting read coverage data were log transformed and normalized for each ChIP sample by dividing by the standard deviation across all 1kb windows and subtracting the 25th percentile across all 1kb windows. For each 1kb window and condition, the resulting data were averaged across replicates. The data were used to calculate the Pearson Correlation coefficient r between all conditions once for autosomes and once for the X chromosome. The distance d = 1 – r was calculated, and hierarchical clustering was used with the complete linkage method to cluster the conditions. The results are displayed in a heatmap where the cell coloring indicates r between two conditions (Fig. S1). The analysis was performed in R version 3.5.1 (R Core Team 2018).

### Metagene plots

Metagene plots for the various ChIP targets and conditions (e.g. Fig. 2C, 4A, and S7) were generated by aligning genes of length greater than 1.25 kb at their TSS and TES using WormBase WS220 gene annotations. Regions 1 kb upstream to 1 kb downstream from the TSS and TES were divided into 150bp windows stepped every 50bp. The mean read coverage within each of these 150bp windows was calculated and normalized for each ChIP data set by dividing by the standard deviation across all 150bp windows and subtracting the 25th percentile across all 150bp windows. For each 150bp window the normalized data were averaged across replicates. A metagene profile for a set of genes was generated by averaging and plotting for each 150bp window the data across the genes in the set. Light vertical lines indicate 95% confidence intervals for the mean of each 150bp window. The analysis was performed in R version 3.5.1.

### Boxplots and scatterplots

To display promoter ChIP signal in boxplots (Fig. S8) and scatter plots (Fig. 4B and S9), the mean read coverage for each protein-coding gene was calculated over the region 250bp up and down from the TSS. For boxplots (Fig. S8) resulting mean read coverage data were log transformed and normalized for each ChIP data set by dividing by the standard deviation across all genes and subtracting the 25th percentile across all genes. For each promoter the resulting signals were averaged across replicates and plotted in Fig S8. In scatterplots (Fig. 4B and S9) the wild-type log2 normalized read coverage was subtracted from the mutant log2 normalized read coverage for each promoter, resulting in a log2 fold change of mutant over wild-type promoter signal.

### Gene set definitions

Ubiquitous genes, originally defined and discussed in (Rechtsteiner *et al.* 2010), are genes that were found to be expressed in germline, muscle, neural, and gut tissues (Wang *et al.* 2009; Meissner *et al.* 2009). Germline-enriched genes are as defined in (Reinke *et* al. 2004). Germline-specific genes are genes whose transcripts were found to be expressed exclusively in the adult germline and maternally loaded into embryos; these genes were defined using multiple datasets as described in (Rechtsteiner *et al.* 2010). Soma-specific genes are genes expressed in at least 1 of 3 somatic tissues (muscle, gut, and/or neuron) with at least 8 SAGE tags (Meissner *et al.* 2009) but not enriched (Reinke *et al.* 2004) or detectably expressed (Wang *et al.* 2009) in the adult germline. Silent genes are 415 serpentine receptor genes that are expressed in a few mature neurons and are not detectably expressed in L1 larvae, originally defined in (Kolasinska-Zwierz *et al.* 2009). HTA germline genes, as defined in (Petrella *et al.* 2011), are genes that were significantly up-regulated in *lin-35(n745)* mutants versus wild type and also significantly down-regulated in *lin-35(n745) mes-4(RNAi)* versus *lin-35(n745),* and that have germline-enriched expression (Reinke *et al.* 2004).

### HTA larval arrest assays

L4 larvae were placed at 26°C for ~18 hours and then moved to new plates and allowed to lay embryos for 8 hours. Progeny were scored for L1 larval arrest (Petrella *et al.,* 2011).

### Immunohistochemistry

L1 larvae were obtained by hatching embryos in the absence of food in M9 buffer and fixed using methanol and acetone (Strome and Wood 1983). Anti-PGL-1 primary antibody (Kawasaki *et al.* 1998) was diluted 1:30,000 and larvae were stained for ~18 hours at 4°C (Petrella *et al.* 2011). Alexa Fluor 488 (Invitrogen) secondary antibody was used at a 1:500 dilution for 2 hours at room temperature. Slides were mounted in Gelutol and imaged using a Nikon Inverted Microscope Eclipse Ti-E confocal microscope at 60X.

## RESULTS

### *lin-15B* mutants lose a large proportion of H3K9me2 promoter peaks; *lin-35* and *lin-37* mutants lose fewer

To better understand how synMuv B proteins regulate germline gene expression in somatic cells, we sought to identify changes in histone modification patterns in mutants compared to wild type. We profiled the distributions of two histone modifications associated with active chromatin (H3K4me3 and H3K36me3) and two histone modifications associated with repressive chromatin (H3K27me3 and H3K9me2) using chromatin immunoprecipitation followed by high-throughput sequencing (ChIP-seq). Experiments were done on L1 animals that experienced embryogenesis at 20°C or 26°C for four genotypes: wild type and three synMuv B mutants, *lin-15B(n744), lin-35(n745),* and *lin-37(n758).* Because L1 stage worms have 550 somatic cells and only 2 germline cells, extracts from L1s contain genomic material primarily from somatic tissue. Analysis of H3K4me3 and H3K36me3 patterns showed increased enrichment of these marks in mutants compared to wild type on classes of genes that are up-regulated in synMuv B mutants (discussed below). As the level of enrichment of these marks generally correlates well with expression level, this change was expected. We saw no changes in the pattern of the repressive modification H3K27me3 between mutants and wild type. However, we observed significant changes in the pattern of the repressive modification H3K9me2 between synMuv B mutants and wild type, especially on germline-expressed genes. We analyzed the changes to H3K9me2 patterns in detail to investigate whether this particular histone modification is important for repression of germline gene expression by synMuv B complexes.

Analysis of H3K9me2 showed there are not genome-wide changes in the distribution of H3K9me2 enrichment on autosomes and the X chromosome between mutants and wild type (Fig. S1). However, a subset of H3K9me2 peaks were observed to be lost or reduced in synMuv B mutants (Fig. 1A and B). To investigate the pattern of this loss/reduction, we performed peak calling for H3K9me2 and designated two types of peaks depending on the location of H3K9me2 relative to coding gene bodies. “Gene body peaks” are those peaks where H3K9me2 overlapped with at least a portion of the coding region of the gene that is more than 250bp downstream of the transcription start site (TSS) (Fig. 1A). The distribution of genes with gene body peaks mirrors what has been previously described for the general pattern of H3K9me2 and H3K9me3 enrichment in the *C. elegans* genome (Fig. 3A; Liu *et al.* 2011; Evans *et al.* 2006). “Promoter peaks” are those peaks where H3K9me2 overlapped with a region 750 bp upstream to 250 bp downstream of the TSS, but not further than 250bp downstream of the TSS (Fig. 1B). In wild-type extracts, H3K9me2 gene body peaks were generally broader than promoter peaks (Fig. 1A and B), and genes with body peaks (2991 at 20°C/2871 at 26°C) were about three times more abundant than genes with promoter peaks (984 at 20°C/981 at 26°C) (Fig. 1C and D).

**Figure 1:**
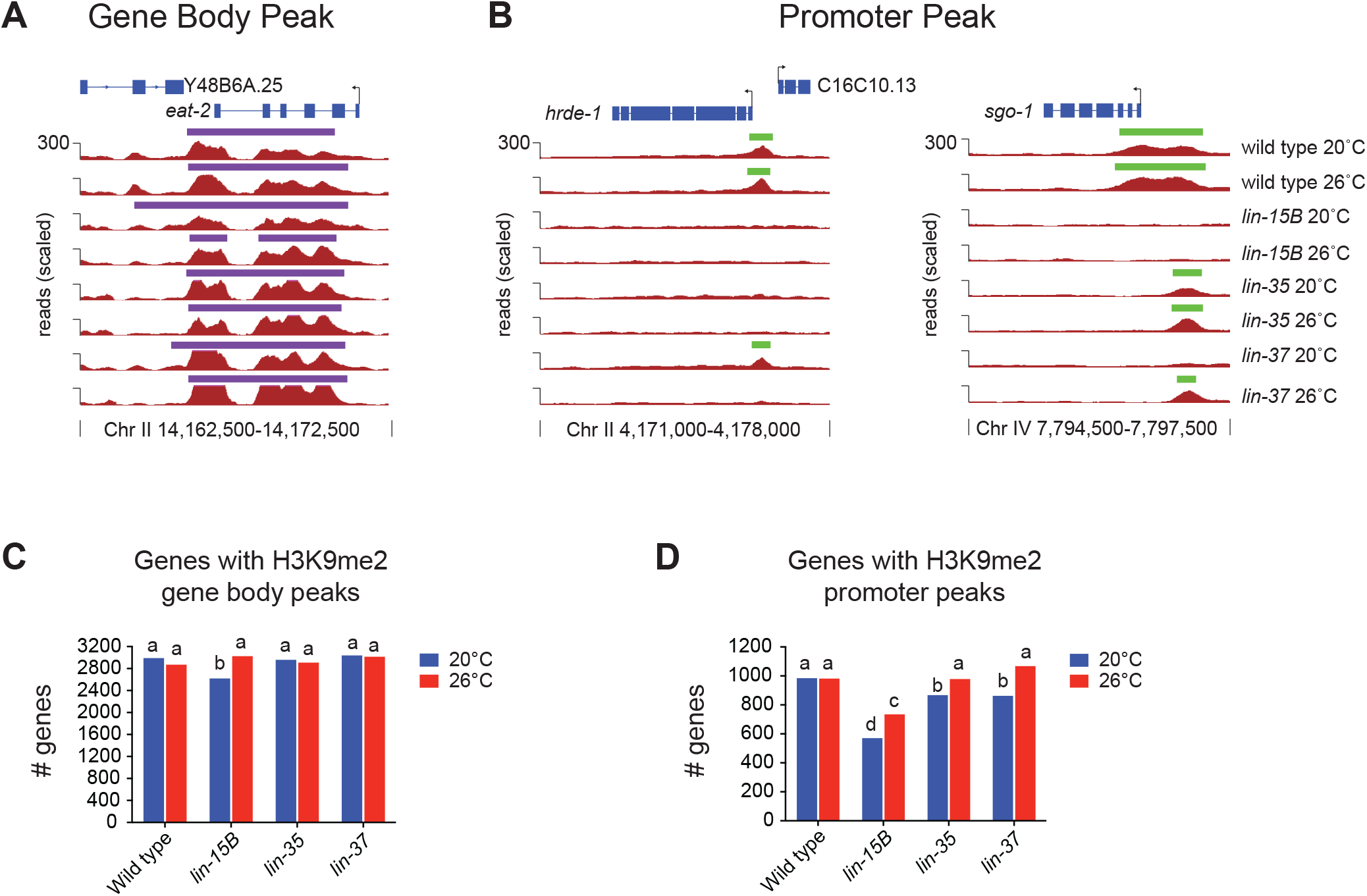
H3K9me2 promoter peaks are lost in *lin-15B* mutant L1s. (A, B) H3K9me2 ChIP-seq data visualized on the UCSC genome browser at one gene *eat-2* (A) with a H3K9me2 gene body peak (peak calls in purple) and at two germline-expressed genes *hrde-1* and *sgo-1* (B) with an H3K9me2 promoter peak (peak calls in green). The vertical lines and arrows indicate the location of the transcription start site (TSS) and the direction of transcription. Signals shown are ChIP-seq reads scaled to 15 million total reads, see Methods. (C) Number of genes in each genotype with a called H3K9me2 peak in the gene body. (D) Number of genes in each genotype with a called H3K9me2 peak in the promoter. Genotypes with statistically significantly different numbers of genes with a called peak are designated with different letters (Chi squared p-value < 0.01).

Our analysis showed that loss of synMuv B proteins had a smaller effect on H3K9me2 in gene bodies than in promoters. In *lin-15B* mutants grown at 20°C ~12% fewer genes had a gene body peak compared to wild type; there was no reduction in the number of genes with H3K9me2 gene body peaks at 26°C (Fig. 1C). In contrast, in *lin-15B* mutants there were significantly fewer genes with H3K9me2 promoter peaks at both 20°C (~42% fewer) and 26°C (~25% fewer) when compared to wild type (Fig. 1D). The H3K9me2 promoter peaks found in *lin-15B* are for the most part a subset of the H3K9me2 promoter peaks in wild type (Fig. S2). Unlike the significant loss of genes with H3K9me2 promoter peaks seen in *lin-15B* mutants, fewer genes with H3K9me2 promoter peaks were lost in *lin-35* and *lin-37* mutants.

In both *lin-35* and *lin-37* mutants, there was no decrease in the number of genes with H3K9me2 gene body peaks (Fig. 1C). There was a small but significant decrease in the number of genes with H3K9me2 promoter peaks compared to wild type at 20°C, but no significant change in the number of genes with promoter peaks at 26°C (Fig. 1D). This is the first description of a molecular difference in phenotypes seen between mutants in DREAM complex members and *lin-15B* mutants and may represent a difference in their molecular function at target loci.

### Genes with an H3K9me2 promoter peak are enriched for DREAM and LIN-15B target genes in wild type but not in *lin-15B* mutants

If localization of H3K9me2 to promoters is driven by synMuv B binding, we predict that genes with H3K9me2 promoter peaks are bound by synMuv B proteins in wild-type animals. There is a high co-occurrence of DREAM complex and LIN-15B binding, with 70% of DREAM bound loci also bound by LIN-15B (Fig. S3). We compared genes with an H3K9me2 promoter peak with genes that we defined as DREAM complex targets and/or LIN-15B targets. We defined 170 DREAM complex targets as those genes bound by the DREAM complex in their promoter by ChIP-seq in late embryos (Goetsch *et al.* 2017) that are also significantly up-regulated in *lin-35* mutant L1s at 26°C (Petrella *et al.* 2011). We defined 115 LIN-15B targets as those genes bound by LIN-15B in their promoter by ChIP-chip in late embryos (this paper) that are also significantly up-regulated in *lin-15B* mutant L1s at 26°C (Petrella *et al.* 2011). Genes with an H3K9me2 promoter peak are enriched for DREAM complex and LIN-15B target genes in wild type, *lin-35,* and *lin-37* mutants but not in *lin-15B* mutants (Fig. 2A). Thus, genes that have H3K9me2 promoter localization are correlated with DREAM complex and LIN-15B binding, and this correlation is disrupted when LIN-15B is absent.

**Figure 2:**
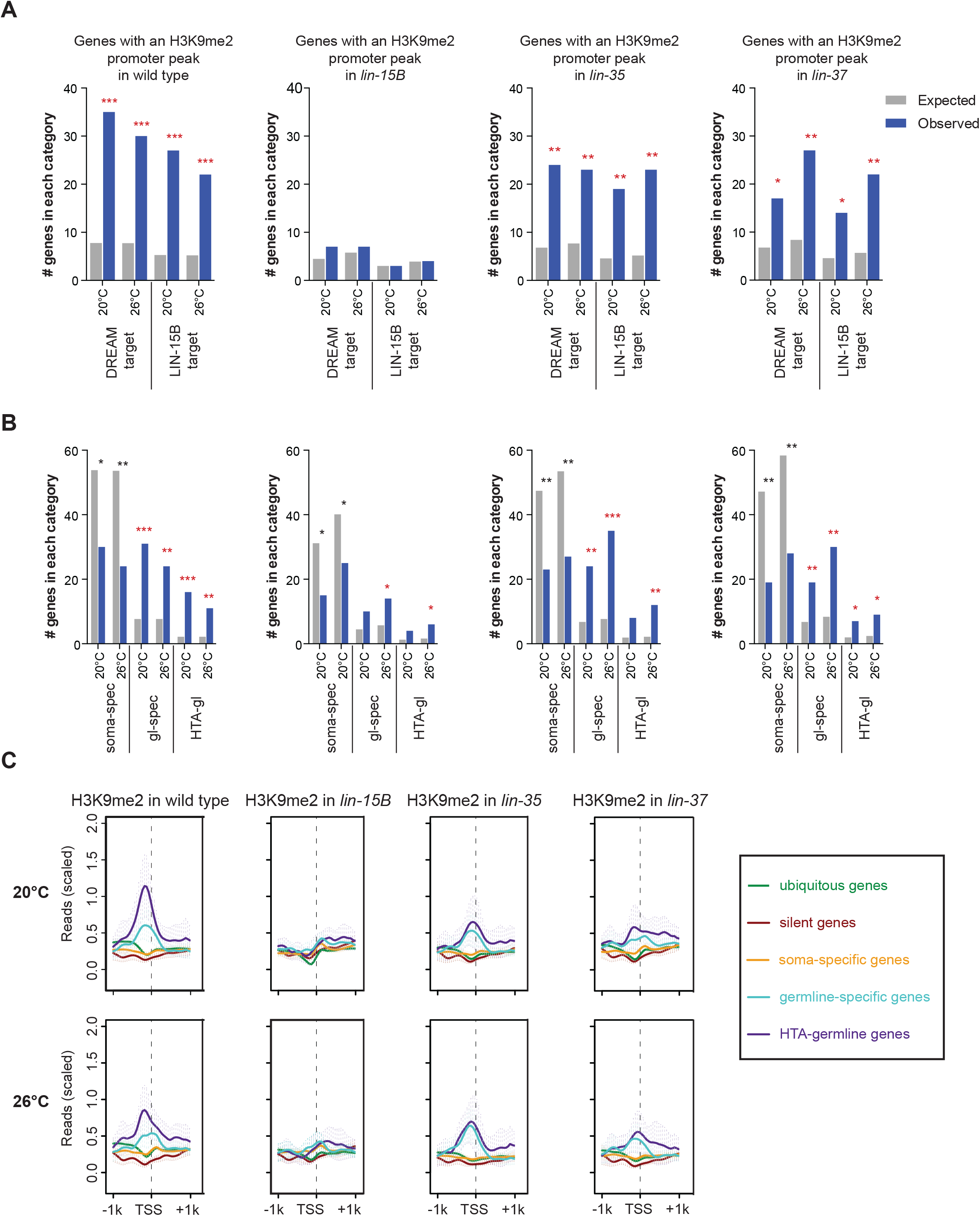
H3K9me2 promoter peaks are associated with synMuv B targets and germline-specific genes in wild-type L1s. (A) Enrichment analysis of genes with an H3K9me2 promoter peak expected by chance and observed among genes that are DREAM complex or LIN-15B targets in the 4 genotypes indicated. DREAM complex targets are defined as genes that are both bound by the DREAM complex in their promoter (Goetsch *et al.* 2017) and also up-regulated in *lin-35* mutants at 26°C (Petrella *et al.* 2011). LiN-15B targets are defined as genes bound in their promoter by LİN-15B (this study) and also up-regulated in *LIN-15B* mutants at 26°C. (B) Enrichment analysis of genes with an H3K9me2 promoter peak expected and observed among genes that are normally expressed specifically in the soma (soma-spec, 1181 genes), expressed specifically in the germline (gl-spec, 169 genes), and genes in the HTA-germline category (HTA-gl, 48 genes) in the 4 genotypes indicated (see Materials and Methods for definitions of gene categories). Significant overenrichment (red) or under-enrichment (black) was determined by the hypergeometric distribution (*p-value < 0.01, **p-value < 1 × 10^−5^, ***p-value < 1 × 10^−10^). (C) Metagene profiles of mean H3K9me2 ChIP-seq signal 1kb up- and downstream from the transcription start site (TSS) for the categories of genes analyzed in B and genes that are normally expressed in all tissues (ubiquitous or ubiq, 2576 genes) and repressed in most tissues (silent, 415 genes). Reads were scaled by dividing by the standard deviation and subtracting the 25th percentile.

### Genes with an H3K9me2 promoter peak lose enrichment for germline genes in *lin-15B* mutants

One of the major phenotypes of many synMuv B mutants, including *lin-15B* mutants, is the ectopic expression in somatic cells of genes whose expression is normally restricted to the germline (Wang *et al.* 2005; Petrella *et al.* 2011; Wu *et al.* 2012). We investigated if genes that have an H3K9me2 promoter peak are enriched for genes that are specifically expressed in the germline. We analyzed four categories of expression: genes that are broadly expressed in all tissues (2576: ubiquitous, ubiq), genes that are repressed in most tissues (415: silent), genes that are expressed specifically in somatic tissues (1181: soma), and genes that are expressed specifically in the germline (169: gl-spec). Genes with H3K9me2 promoter peaks in wild type are enriched for genes with germline-specific expression, but not for genes with ubiquitous, silent, or somatic expression (Fig. 2B and S4). These enrichments are mirrored when plotting H3K9me2 ChIP-seq signal around the transcription start site (TSS) averaged over the genes in each expression category (Fig. 2C). If H3K9me2 at germline gene promoters is correlated with synMuv B repression of germline gene expression in the soma, then we would predict that germline genes would lose H3K9me2 promoter peaks in synMuv B mutants. Indeed, in *lin-15B* mutants, there were many fewer germline-specific genes with an H3K9me2 promoter peak, and there was a large decrease in the signal of H3K9me2 at the TSS of germline-specific genes (Fig. 2B and 2C). *lin-35* and *lin-37* mutants resembled wild type in showing genes with an H3K9me2 promoter peak enriched for germline-specific genes (Fig. 2B and 2C).

We also examined germline genes whose misregulation is correlated with the high temperature larval arrest (HTA) phenotype (Petrella *et al.* 2011). HTA-germline targets are defined as genes normally expressed in the germline that are up-regulated in arrested *lin-35* mutant L1s at 26°C and whose expression returns to near wild-type levels in HTA-suppressed *lin-35; mes-4(RNAi)* double mutant L1s at 26°C (48: HTA-gl) (Petrella *et al.* 2011). Similar to what was seen with germline-specific genes, genes with an H3K9me2 promoter peak were enriched for HTA-germline genes in wild type, *lin-35,* and *lin-37* mutants, but this enrichment was reduced in *lin-15B* mutants (Fig. 2B). These data together reveal a striking loss of H3K9me2 over the promoters of germline-specific and HTA-germline genes in *lin-15B* mutants, but not *lin-35* or *lin-37* mutants.

### H3K9me2 promoter peaks are distributed along the length of autosomes

Previous work on H3K9me2 in *C. elegans* focused on its distribution in broad domains on autosomal arms and the role of H3K9me2 in repressing repetitive sequences (Ikegami *et al.* 2010; Liu *et al.* 2011; Guo *et al.* 2015; Zeller *et al.* 2016). Little investigation has been done into what role the more narrowly focused H3K9me2 found in promoters may be serving in gene regulation. In *C. elegans,* genes with expression that is higher in the germline than other tissues (germline-enriched genes) or with expression exclusive to the germline (germline-specific genes), show a biased localization to the centers of autosomes compared to the localization of all coding genes (Fig. S5). Therefore, if H3K9me2 promoter peaks are associated with regulation of germline gene expression, we would predict that H3K9me2 promoter peaks would also be found in the center regions of chromosomes and not be biased to the arms. We compared the distributions along autosomes of genes with H3K9me2 in their gene body versus in their promoter. In wild type, genes with H3K9me2 in their gene body demonstrated the previously reported pattern of H3K9me2 enrichment on autosomal arms compared to centers (Fig. 3A). For genes with an H3K9me2 gene body peak, all mutants showed the same autosomal arm bias as seen in wild type (Fig. 3A, 3B, and S6). In contrast, genes with an H3K9me2 promoter peak in wild type were more evenly distributed across autosomes, with weak or no depletion from autosomal centers (Fig. 3A and B). Notably, *lin-15B* mutants showed strong depletion of genes with H3K9me2 in their promoter on all autosomal centers (Fig. 3A and B), suggesting that LIN-15B is needed for H3K9me2 localization on gene promoters in autosomal centers where germline genes are enriched. *lin-35* and *lin-37* mutants showed distributions of genes with H3K9me2 in their promoter similar to wild type (Fig. S6). H3K9me2 promoter peaks in chromosome centers in wild type represent a pattern not previously described for H3K9me2 in *C. elegans* and place H3K9me2 promoter peaks in mainly euchromatic regions where they may affect coding gene expression. Additionally, the loss of H3K9me2 from promoter peaks in autosomal centers in *lin-15B* mutants suggests that LIN-15B plays a specific role in directing H3K9me2 to areas of the genome where there are fewer repeats and more coding genes, especially germline genes.

**Figure 3:**
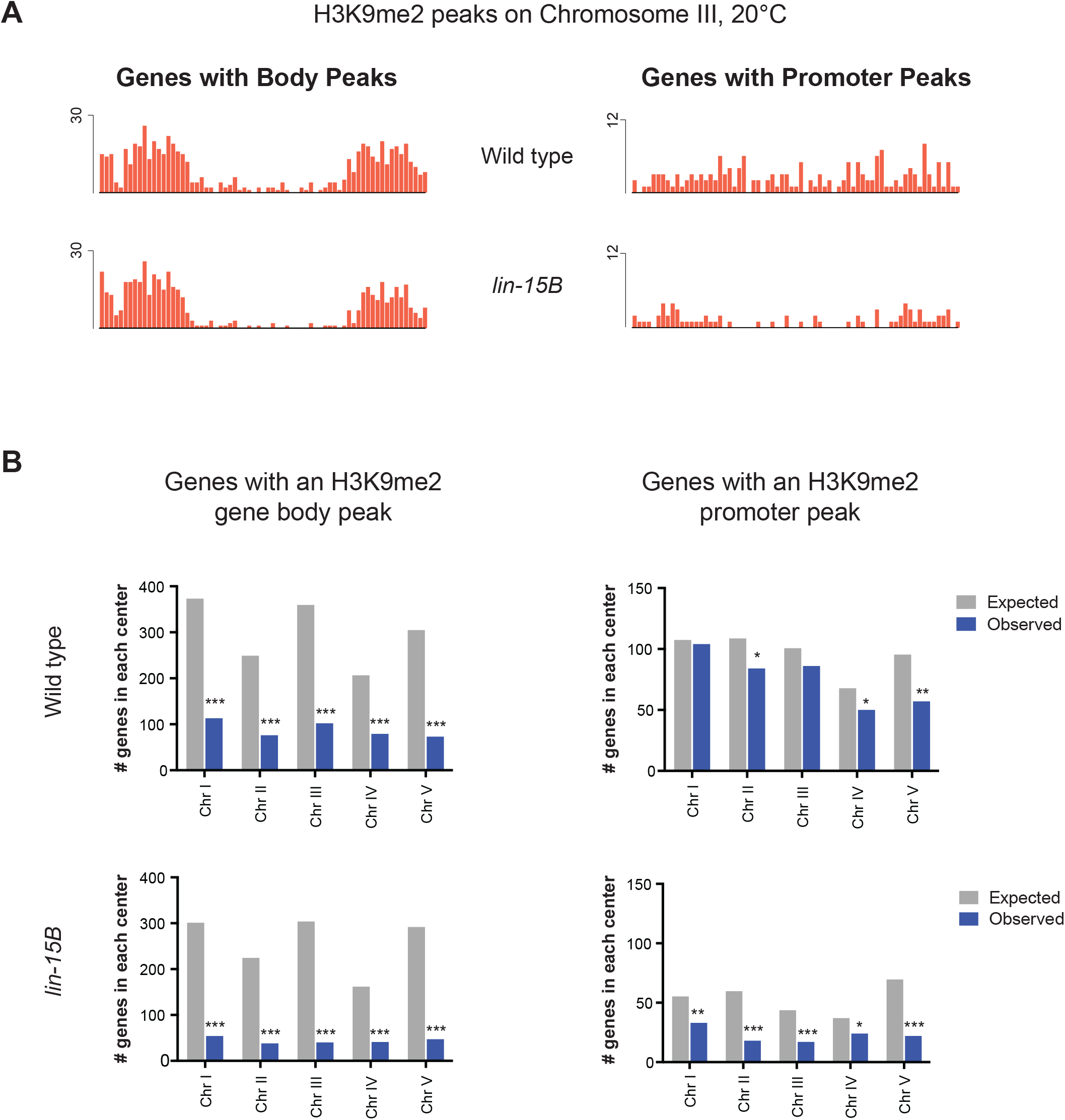
Genes with an H3K9me2 promoter peak in wild-type L1s are not biased toward autosomal arms. (A) Binned distribution of genes with an H3K9me2 gene body or promoter peak at 20°C in 200 kb windows across chromosome III in wild-type and *lin-15B* mutant L1s. (B) Enrichment analysis of genes with an H3K9me2 gene-body peak or promoter peak expected by chance and observed in chromosome centers in wild-type and *lin-15B* mutant L1s. The expected number is based on the percentage of coding genes in the center versus arm regions of each chromosome; the observed number is the number of genes in the chromosome centers at 20°C. The locations of chromosome arm and center boundaries are from (Liu *et al.* 2010). Significant under-enrichment (black) was determined by the hypergeometric distribution (*p-value < 0.01, **p-value < 1 × 10^−5^, ***p-value < 1 × 10^−10^). Error bars indicate 95% confidence intervals for the mean (also see Materials and Methods).

### Loss of H3K9me2 in mutants is associated with increased H3K4me3 on germline genes

Trimethylation of histone H3 on lysine 4 (H3K4me3) and lysine 36 (H3K36me3) are correlated with active gene expression (Liu *et al.* 2011; Ho *et al.* 2014; Evans *et al.* 2016). Thus, we expected to see increases in H3K4me3 and H3K36me3 on germline genes in synMuv B mutants. Indeed, synMuv B mutants displayed increases in H3K4me3 and H3K36me3 on germline-specific and HTA-germline but not on other categories of genes (Fig. 4A, S7, and S8). The increased levels of both H3K4me3 and H3K36me3 on germline-specific and HTA-germline genes in mutants is consistent with these genes being expressed at higher levels, most likely in a larger population of cells (i.e. somatic cells in addition to the 2 primordial germ cells) in these mutants.

**Figure 4:**
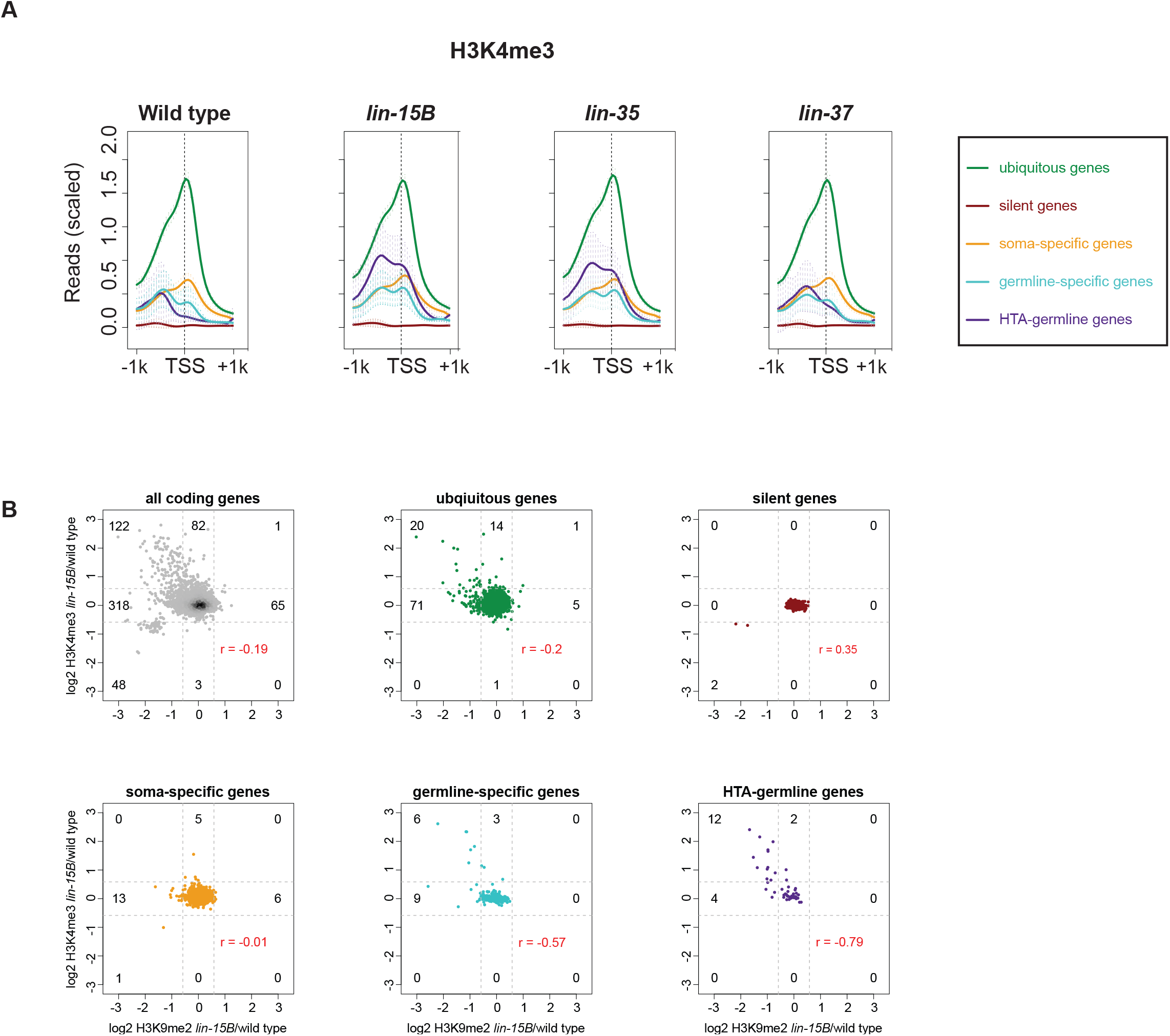
H3K4me3 increases on germline genes that lose H3K9me2. (A) Metagene profiles of mean H3K4me3 ChIP-seq signal 1kb up- and downstream from the transcription start site (TSS) for genes that show ubiquitous, silent, soma-specific, germline-specific, or HTA-germline expression (see Materials and Methods for definitions of gene categories). Reads were scaled by dividing by the standard deviation and subtracting the 25th percentile. Error bars indicate 95% confidence intervals for the mean. (B) Scatter plots of log2 fold change of the H3K9me2 signal over the TSS in *lin-15B* mutant/wild type vs. log2 fold change of the H3K4me3 signal over the TSS in *lin-15B* mutant/wild type. The signal was calculated within 250bp upstream and downstream of the transcription start site (TSS) at 20°C. Shown are all coding genes and genes with ubiquitous, silent, soma-specific, germline-specific, or HTA-germline expression. Dotted lines represent 1.5 fold cut-offs; the numbers of genes above and below the cut-offs are indicated. r values show the Pearson correlation between changes in H3K9me2 and changes in H3K4me3 for each set of genes.

Because loss of H3K9me2 may not be sufficient to lead to gene expression changes, we investigated if there is a correlation between loss of the repressive H3K9me2 chromatin modification and gain of the active H3K4me3 chromatin modification in gene promoters. To compare those marks in promoter regions, we calculated the log2 fold change of the signal of each modification in *lin-15B* mutant/wild type within 250bp upstream and downstream of the transcription start site (TSS). A higher histone modification signal in *lin-15B* mutants than wild type would result in a positive log2 fold change; a lower histone modification signal in *lin-15B* mutants than wild type would result in a negative log2 fold change. In *lin-15B* mutants, 25% of all genes (122 of 448) that had reduced H3K9me2 promoter signal also had increased H3K4me3 promoter signal, using a 1.5-fold cut-off (Fig. 4B). Strikingly, 40% of germline-specific (6 of 15) and 75% of HTA-germline genes (12 of 16) that had reduced H3K9me2 promoter signal also had increased H3K4me3 promoter signal (Fig. 4B). Thus, in *lin-15B* mutants loss of H3K9me2 enrichment on the promoter of germline genes is more likely to result in increased H3K4me3 promoter enrichment than on genes in other categories. Interestingly, although fewer genes showed a decrease in promoter H3K9me2 enrichment in *lin-35* and *lin-37* mutants, the percentages of germline-specific and HTA-germline genes that showed reduced promoter H3K9me2 and increased promoter H3K4me3 were similar to the percentages in *lin-15B* mutants (Fig. S9). Thus, the germline genes that lose H3K9me2 promoter enrichment in any of the three mutants have a correlated increased enrichment of H3K4me3 and up-regulation in synMuv B mutants.

### Global loss of H3K9me2 leads to phenotypes similar to *lin-15B* and DREAM complex mutants

To investigate if loss of H3K9me2 promoter localization plays an important role in *lin-15B* mutant phenotypes, we analyzed mutants for the histone methyltransferases (HMTs) responsible for H3K9 methylation. Loss of these HMTs leads to a global loss of all H3K9 methylation, which may result in a phenocopy of *lin-15B* mutants. H3K9 methylation in *C. elegans* embryos is catalyzed by two HMTs, MET-2 and SET-25, which primarily catalyze H3K9me1/2 and H3K9me3, respectively (Towbin *et al.* 2012). If loss of H3K9 methylation is associated with ectopic germline gene expression and the HTA phenotype, we would expect that *met-2* and *set-25* mutants would show these phenotypes. *set-25* single mutants, which lose H3K9me3, showed neither an HTA phenotype nor an ectopic germline gene expression phenotype, as assessed by staining for the germline-specific protein PGL-1 (Fig. 5A and B).

**Figure 5:**
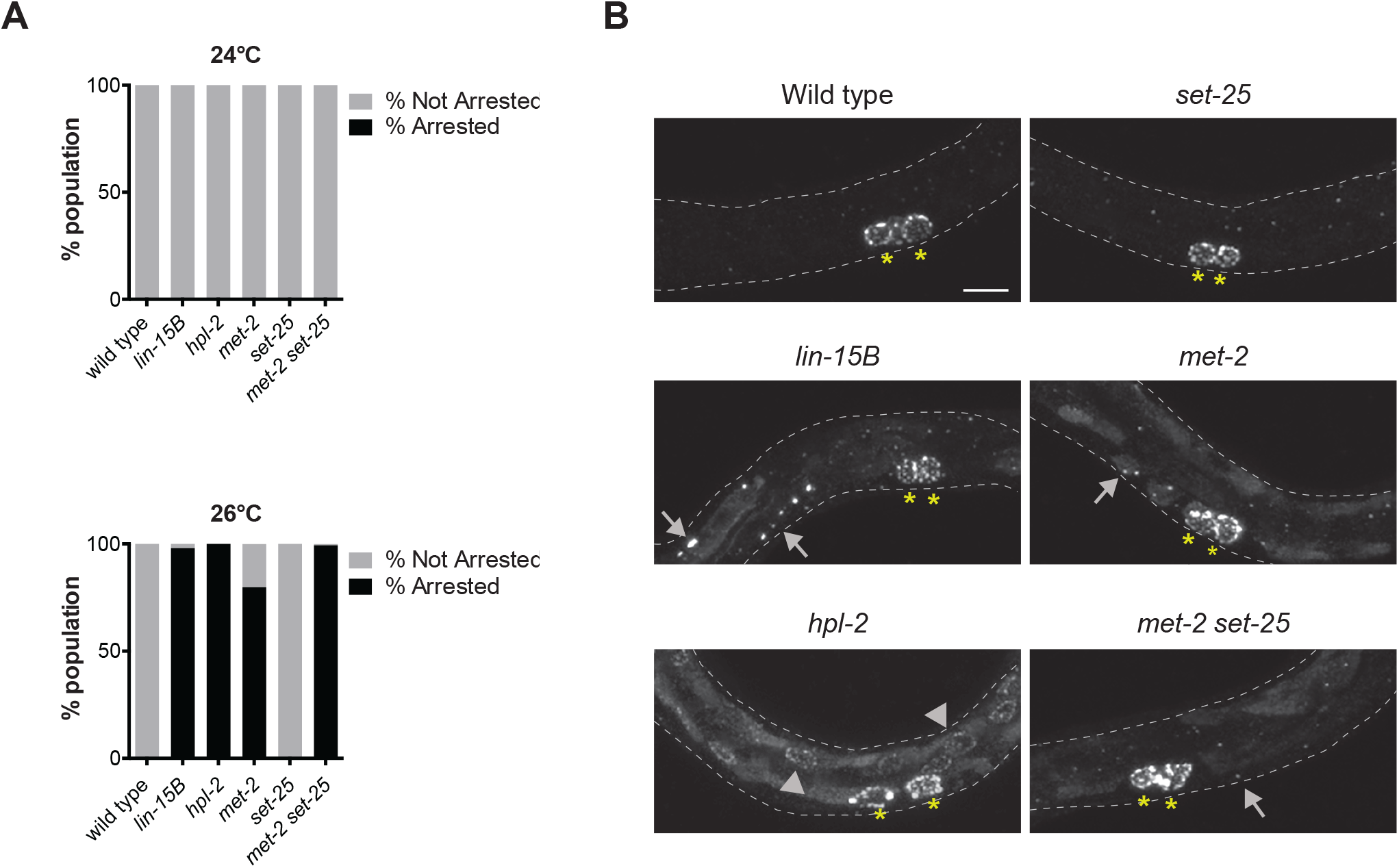
Complete loss of H3K9me2 during development phenocopies synMuv B mutants. (A) The percentage of F1 animals that arrested at a larval stage before L4 was assessed for all genotypes indicated after parent hermaphrodites were upshifted from 20°C to 24°C or 26°C at the L4 stage. (B) Assessment of ectopic expression of PGL-1 in L1 animals at 26°C. Yellow asterisks indicate the two primordial germ cells in which PGL-1 is solely expressed in wild type. Arrowheads indicate ectopic perinuclear punctate PGL-1 in intestinal cells. Arrows indicate ectopic punctate PGL-1 that is not perinuclear. Scale bar: 10μm.

Therefore, H3K9me3 does not appear to be important for repression of germline genes in the soma. In contrast, *met-2* single mutants, which lose 80-90% of H3K9me2 and ~70% of H3K9me3 (Towbin *et al.* 2012), displayed ~80% larval arrest around the L3 stage at 26°C, but no larval arrest at 24°C (Fig. 5A). Thus, *met-2* mutants show an HTA phenotype similar to but weaker than *lin-15B* mutants (Fig. 5A; Petrella *et al.* 2011). We also observed ectopic expression of PGL-1 in *met-2* mutants at 26°C similar in level and distribution to *lin-15B* mutants, with a combination of diffuse and punctate cytoplasmic PGL-1 in intestinal cells (Fig. 5B). Unlike the previously published pattern of ectopic PGL-1 expression in *met-2* (Wu *et al.* 2012), we did not see perinuclear punctae in intestinal cells (Fig. 5B). Perinuclear PGL-1 in intestinal cells has been described for *hpl-2* mutants, the worm homolog of HP1, which is known to bind H3K9me2 (Petrella *et al.* 2011; Wu *et al.* 2012). Thus, *met-2* single mutants have phenotypes similar to, but in the case of HTA weaker, *lin-15B* mutants. To test if the remaining 10-20% of H3K9me2 catalyzed by SET-25 in *met-2* mutants (Towbin *et al.* 2012) partially represses germline gene expression in somatic cells, we analyzed *met-2 set-25* double mutants, which have been shown to completely lack H3K9 methylation during embryonic stages (Towbin *et al.* 2012; Garrigues *et al.* 2015). Consistent with residual SET-25-mediated H3K9me2 serving a role in repression of germline genes in somatic cells, *met-2 set-25* double mutants showed significantly enhanced larval arrest at 26°C when compared to *met-2* single mutants (Fig. 5A). Similar to *lin-15B, lin-35,* and *lin-37* mutants, *met-2 set-25* double mutants did not show increased larval arrest at 24°C. The level and distribution of ectopic PGL-1 in *met-2 set-25* double mutants at 26°C were similar to those seen in *met-2* single mutants (Fig. 5B). Altogether, our results show that a global loss of H3K9me2 phenocopies both the HTA and ectopic germline gene expression seen in synMuv B mutants.

## Discussion

Repression of germline gene expression in the soma is vital, as loss of germline gene repression is a hallmark of various disease states including cancer. Investigating the changes to chromatin that occur when germline genes are misexpressed in the somatic cells of mutants is a first step in understanding the mechanisms that repress germline genes to protect somatic fate and development. Here we investigated the changes to histone modifications that occur in a subset of synMuv B mutants that misexpress germline genes in the soma. We defined a new localization for the repressive histone modification H3K9me2 in wild-type extracts, at the promoter of coding genes, which unlike the previously described broad domains of H3K9me2 are not enriched on autosomal arms (Liu *et al.* 2011; Garrigues *et al.* 2015; Evans *et al.* 2016; Ahringer and Gasser 2018). Promoter enrichment of H3K9me2 in autosomal centers provides a new regulatory role for H3K9me2 in addition to its well described regulation of repetitive elements on autosomal arms. We also found that in wild-type somatic cells genes with an H3K9me2 promoter peak are enriched for genes expressed specifically in the germline and genes that are synMuv B targets. The localization of H3K9me2 to germline genes and synMuv B targets is disrupted strongly in *lin-15B* mutants and weakly in DREAM complex mutants. We additionally showed that loss of H3K9me2 but not H3K9me3 phenocopies synMuv B mutants. Our data implicate H3K9me2 promoter enrichment as an important aspect of repression of germline gene expression in somatic cells.

There is strong evidence that a memory of gene expression/repression and associated chromatin modifications are transmitted from the parental germline to the developing embryo (Furuhashi *et al.* 2010; Rechtsteiner *et al.* 2010; Zenk *et al.* 2017; Tabuchi *et al.* 2018). For example, genes that were expressed in the germline continue to be marked with MES-4-generated H3K36me3 in embryos, even in the absence of ongoing transcription in embryos (Furuhashi *et al.* 2010; Rechtsteiner *et al.* 2010; Kreher *et al.* 2018). It is thought that H3K36me3 marks these genes for re-expression in the germline during post-embryonic development. How then are germline genes repressed properly in somatic tissues when those tissues inherit marks of active expression that potentially set germline genes up for reexpression? Our data, along with other recent work, strongly implicate deposition of H3K9me2 at the proper time in development in creating proper expression patterns in differentiating somatic cells. Recent experiments have shown that in *C. elegans* H3K9me2 and H3K9me3 levels are very low in the nuclei of early stage embryos and only start to accumulate when cells are transitioning from early embryogenesis to mid-embryogenesis (Mutlu *et al.* 2018). This is in part driven by the nuclear import of an active MET-2 complex that catalyzes conversion of H3K9me1 to H3K9me2. The timing of MET-2 import just precedes the stage in embryogenesis when zygotic transcription is up-regulated and when tissue-specific expression patterns emerge (Spencer *et al.* 2011; Levin *et al.* 2012; Robertson and Lin 2015; Mutlu *et al.* 2018). Our data suggest that loss of H3K9me2, either through loss of the MET-2 and SET-25 HMTs that catalyze the mark or through loss of proper localization of H3K9me2 to germline genes in *lin-15B* mutants, leads to misexpression of germline-specific genes in somatic cells. We hypothesize that specific localization of H3K9me2 to germline gene promoters facilitated by LIN-15B is an important aspect of resetting the chromatin landscape of germline genes at this stage of development to prevent their expression in somatic lineages.

A striking aspect of our findings is the difference in changes to promoter-enriched H3K9me2 between *lin-15B* mutants and DREAM complex mutants. It was previously proposed, based on phenotype analysis, that LIN-15B is a member of the DREAM complex (Wu *et al.* 2012). Our data indicate that, although LIN-15B binds to and represses many of the same genes as the DREAM complex, its molecular function at those genes is probably distinct. The proposed DNA-binding domain of LIN-15B may allow it to be independently recruited to similar targets as the DREAM complex, where the two may function together to repress genes. This scenario has implications for regulation of gene expression in the germ line as well as in the soma. Recent work from the Seydoux lab has further implicated the differential presence of LIN-15B in the germline versus soma in this regulation (Lee *et al.* 2017). Maternally provided LIN-15B is normally removed from the primordial germ cells (PGCs), while DREAM components are not (Lee *et al.* 2017). Our work suggests that loss of LIN-15B from the PGCs may protect essential germline genes from being H3K9 methylated and repressed in those cells. How the different synMuv B complexes work together to fully repress germline-specific genes in somatic cells is still an open question. The establishment of H3K9me2 may be an initiating step in germline gene repression or may be one aspect of a series of redundant steps necessary to repress germline genes. Analysis of the order and dependency of DREAM, LIN-15B, and MET-2 binding to germline genes is necessary to address these questions.

The work presented here focuses on a subset of germline-promoting genes that are regulated through the LIN-15B/H3K9me2/DREAM complex pathway. Although this pathway may only regulate a small subset of genes in this way, the repercussions to development are clear, in that the organisms cannot thrive in the face of challenges (e.g. high temperature) when these fate changes occur. Recent work in *Drosophila* underscores the importance of H3K9 methylation in repression of a small subset of coding genes to maintain proper cell fate. In the *Drosophila* ovary, loss of H3K9me3 leads to up-regulation of testis-specific transcripts that change the fate of ovarian germ cells, leading to sterility (Smolko *et al.* 2018). As in *C. elegans*, prior investigations of H3K9 methylation loss in *Drosophila* had focused primarily on up-regulation of repetitive elements (Rangan *et al.* 2011; Wang *et al.* 2011; Guo *et al.* 2015; Zeller *et al.* 2016). However, it is clear that H3K9me2/3 loss leading to up-regulation of small sets of coding genes in a tissue-specific manner can have profound effects on cellular fate and function. As more studies investigate the roles of H3K9me2/3 in repression of coding genes, it seems likely that new pathways will be uncovered that are necessary to create different patterns of H3K9me2/3 in different tissues for maintenance of proper cell fate.

The expression of germline-specific genes in somatic tissues leads to a variety of adverse consequences in diverse animal species. These include L1 starvation and reduced apoptosis during development in *C. elegans* synMuv B mutants, tumor formation in *Drosophila l(3)mbt* mutants, and poor outcomes in human tumors that express germline genes (Janic *et al.* 2010; Petrella *et al.* 2011; Whitehurst 2014; Al-Amin *et al.* 2016). Thus, there is a need across species to repress germline gene expression in the soma to facilitate proper development and somatic function. Our data suggest that repression of germline genes during development in somatic tissues through H3K9me2 may be a conserved mechanism. As in *C. elegans* embryonic somatic cells, mammalian ES cells also repress expression of germline genes (Blaschke *et al.* 2013). Mouse ES cells have been shown to lose repression of germline genes when treated with vitamin C (Blaschke *et al.* 2013; Ebata *et al.* 2017). Interestingly, ectopic expression of germline genes upon exposure of ES cells to vitamin C is dependent on loss of H3K9me2 and DNA methylation. Loss of H3K9me2 at germline genes in response to vitamin C appears to be through the activity of H3K9 demethylases (Ebata *et al.* 2017). In contrast, in *C. elegans* synMuv B mutants, germline genes likely lose H3K9me2 due to failure in the initial HMT-catalyzed deposition of H3K9me2 during development. The conservation of H3K9me2 on germline genes and its role in repressing these genes in developing somatic lineages may represent an ancient regulatory role for H3K9me2. Since in both *C. elegans* and *Drosophila* repression of germline genes in the soma is through complexes known to interact with chromatin (Janic *et al.* 2010; Petrella *et al.* 2011; Wu *et al.* 2012), it will be interesting to see if ectopic expression of germline genes in human somatic tumors is due to loss of these conserved complexes. Finally, not all germline genes, but only a specific subset, are ectopically expressed in these models. Why only certain germline genes are vulnerable to misexpression, if those genes are the same across species, and which cellular processes are disrupted as a result of germline gene misexpression singularly or as a group, are open questions. Further investigation into these questions could have broad implications for understanding conserved basic chromatin mechanisms and therapeutic targets for cancer treatment.

## ACKNOLEDGMENTS

Many thanks to Anita Manogaran for comments and discussion of the manuscript.

Some strains were provided by the CGC, which is funded by NIH Office of Research Infrastructure Programs (P40 OD010440). This work was supported by a NIH grants R00GM98436 and R15GM122005 to L.N.P and NIH grant R01GM34059 to S.S.

---

**Figure S1:**
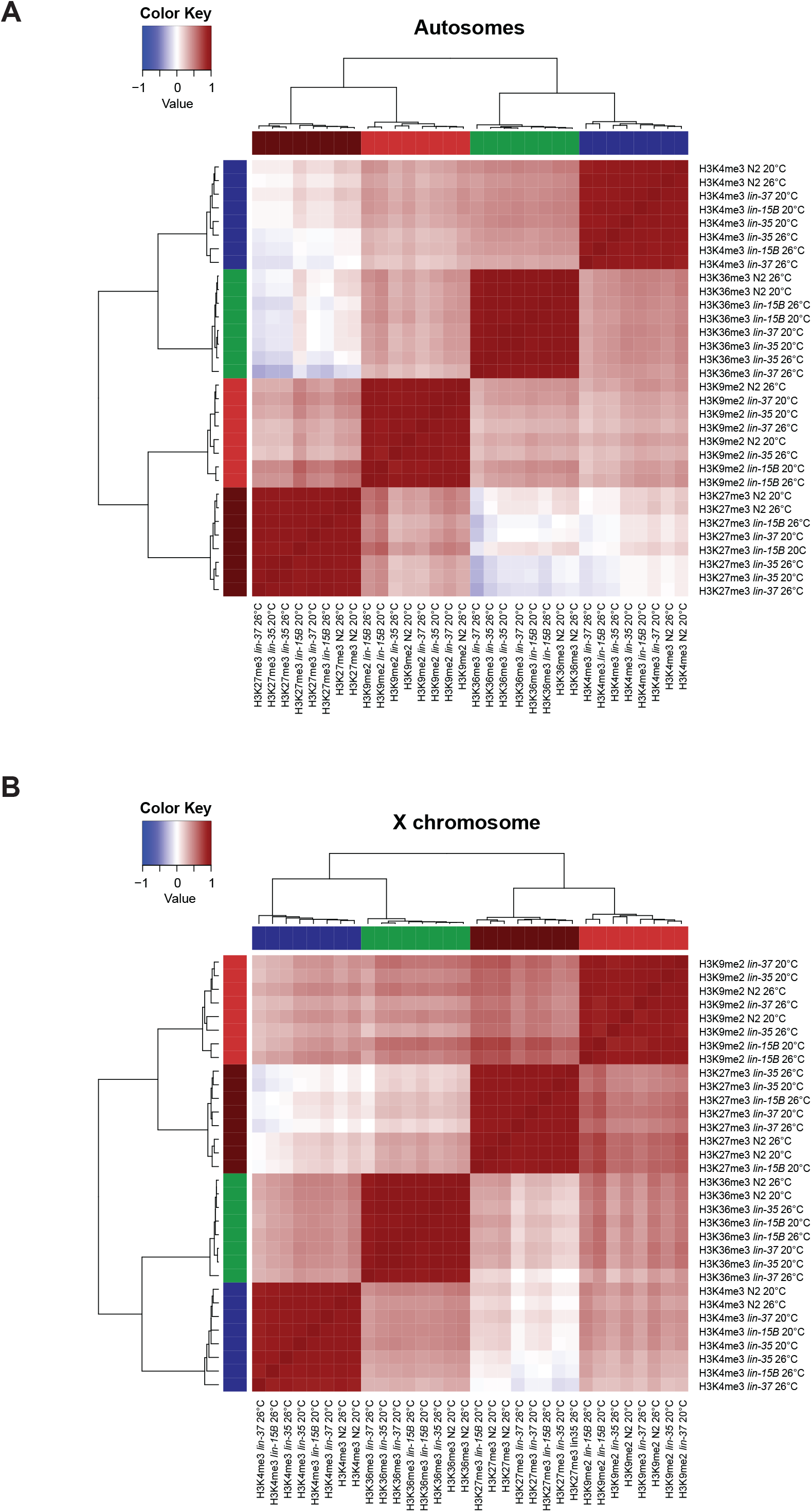
Heatmap and clustering of genome-wide ChIP-seq signals in genotypes and temperatures based on Pearson correlation. (A,B) Heatmap and hierarchical clustering of H3K9me2, H3K4me3, and H3K36me3 ChIP-seq signals in 1kb windows in the different genotypes and temperatures based on Pearson correlation on the autosomes (A) and the X chromosome (B).

**Figure S2:**
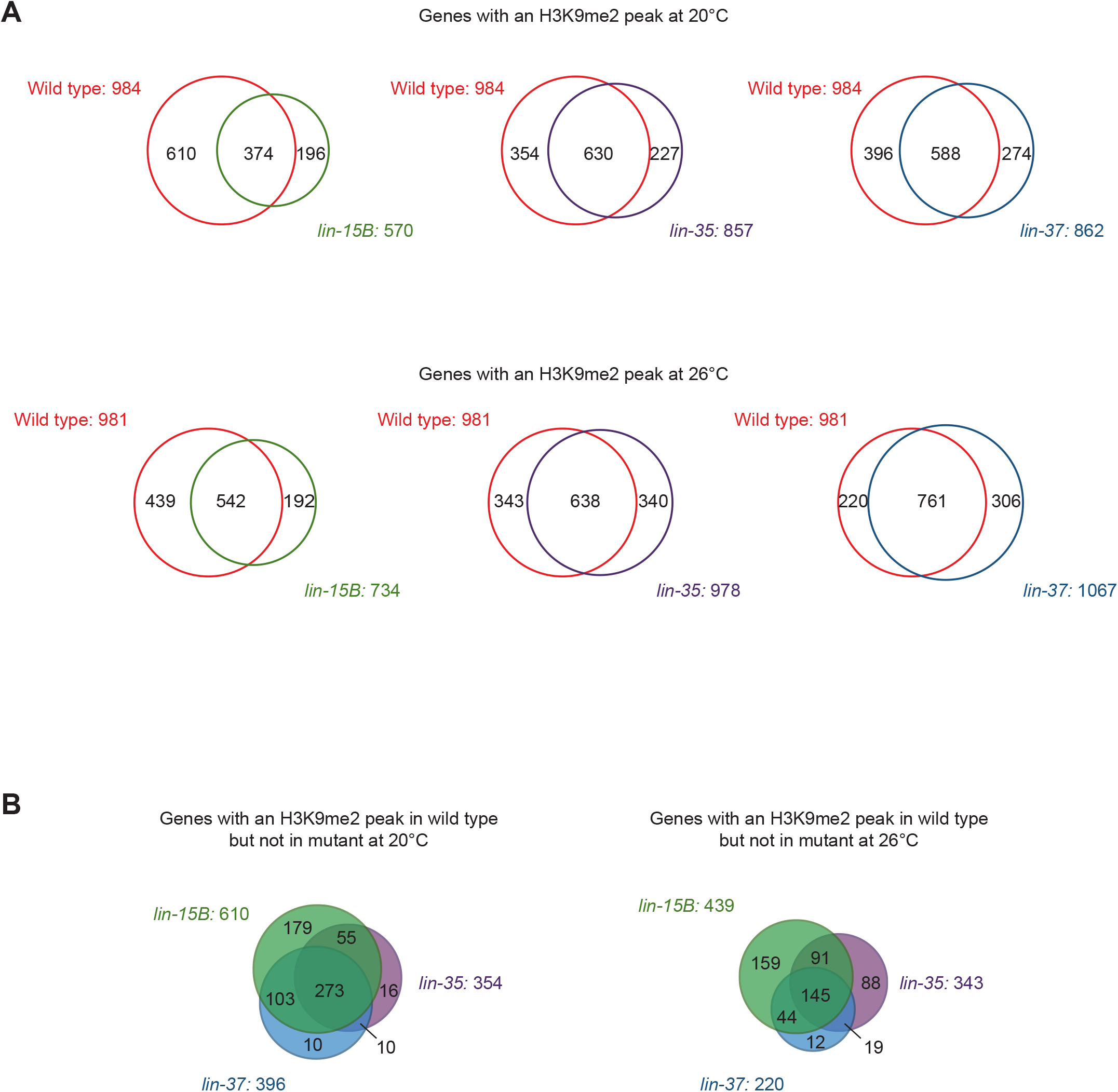
Overlaps of genes with called H3K9me2 promoter peaks amongst different genotypes. (A) Venn diagrams of genes with a called H3K9me2 promoter peak in wild type and each mutant at either 20°C or 26°C. (B) Venn diagrams of genes that lost a called H3K9me2 promoter peak in each mutant at either 20°C or 26°C.

**Figure S3:**
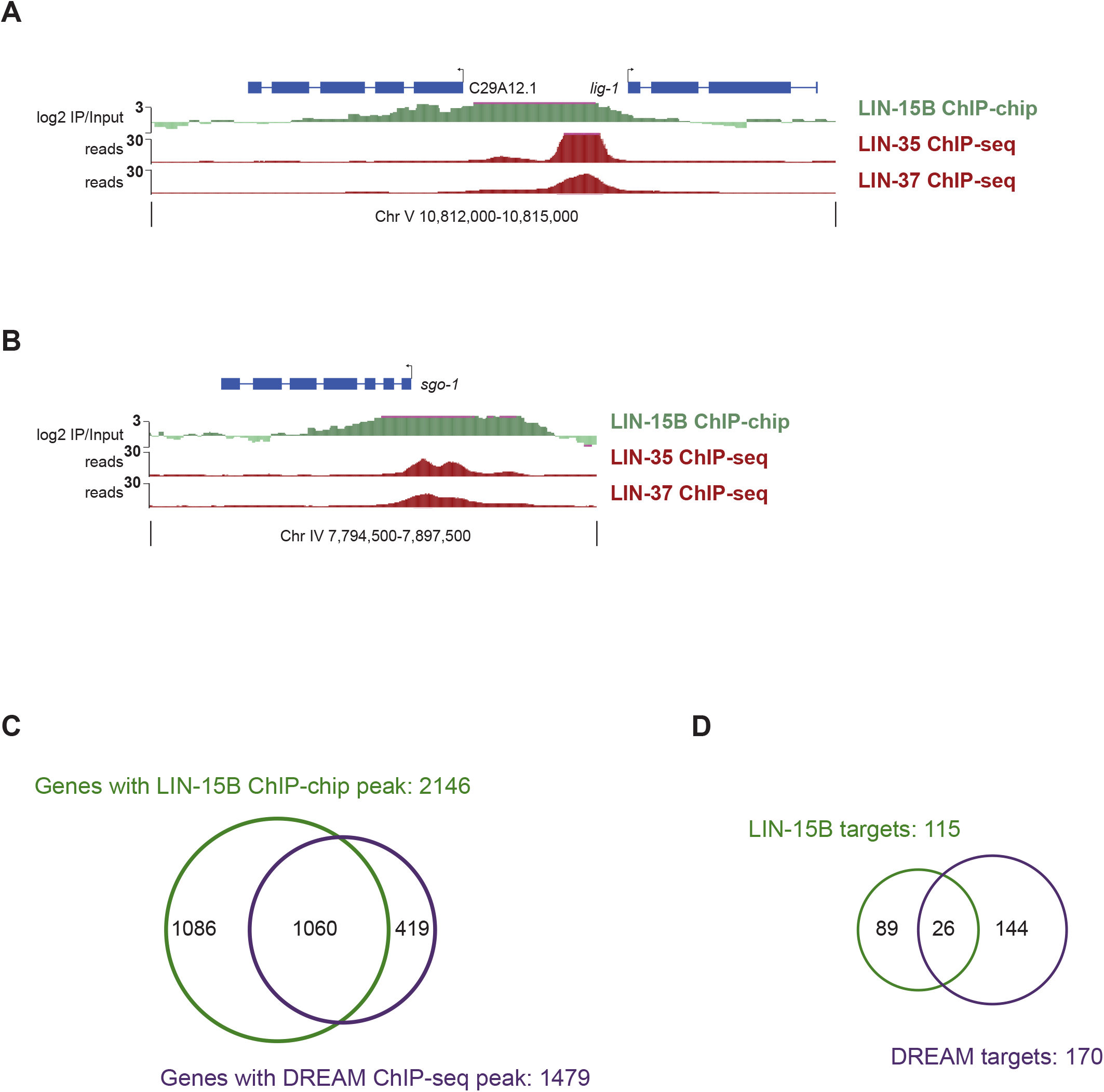
Overlap of LIN-15B and DREAM complex ChIP data. (A, B) LIN-15B (green), LIN-35 (red), and LIN-37 (red) ChIP data visualized on the UCSC Genome Browser at two bidirectional genes C29A12.1 and *lig-1* (A) and the *sgo-1* gene (B) with all three proteins in their promoters. The vertical lines and arrows indicate the location of the transcription start site (TSS) and the direction of transcription. Signals shown are LIN-15B ChIP-chip z-scored log2 fold changes IP/Input and LIN-35 and LIN-37 ChIP-seq reads scaled to 15 million total reads (see Materials and Methods). Note that the broader LIN-15B peaks are at least partially expected due to the different ChIP technologies (ChIP-seq generally has more dynamic range than array based ChIP-chip) and the log transformation of the ChIP-chip ratios. (C) Venn diagram of genes with a called LIN-15B promoter peak and DREAM promoter peak (Goetsch *et al.* 2017). (D) Venn diagram of genes defined as LIN-15B targets and DREAM complex targets.

**Figure S4:**
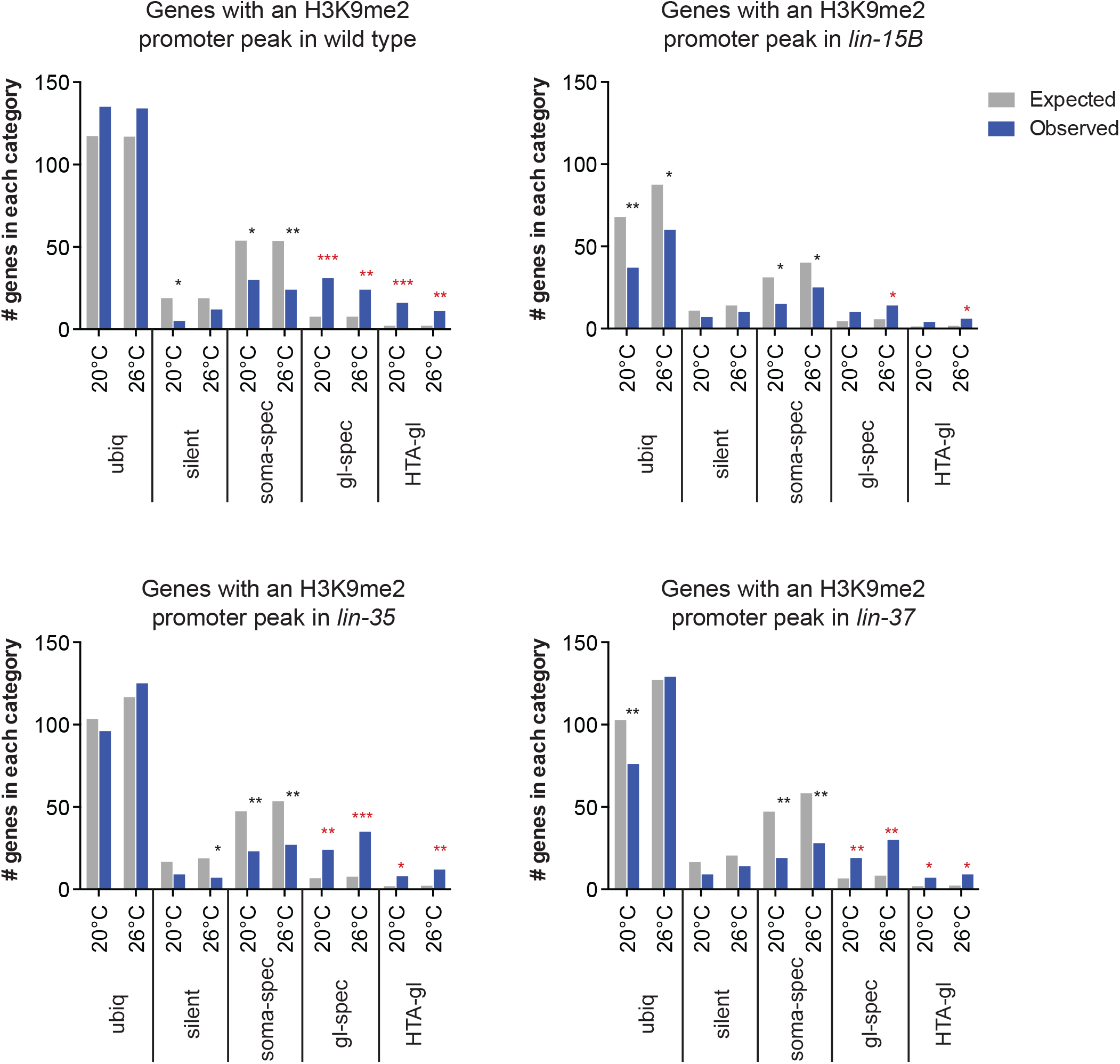
Enrichment analysis of genes with called H3K9me2 promoter peaks for different categories of expression. Enrichment analysis of genes with an H3K9me2 promoter peak expected and observed among genes that show ubiquitous, silent, soma-specific, germline-specific, or HTA-germline expression. Significant over-enrichment (red) or under-enrichment (black) was determined by the hypergeometric distribution (*p-value < 0.01, **p-value < 1 × 10^−5^, ***p-value < 1 × 10^−10^).

**Figure S5:**
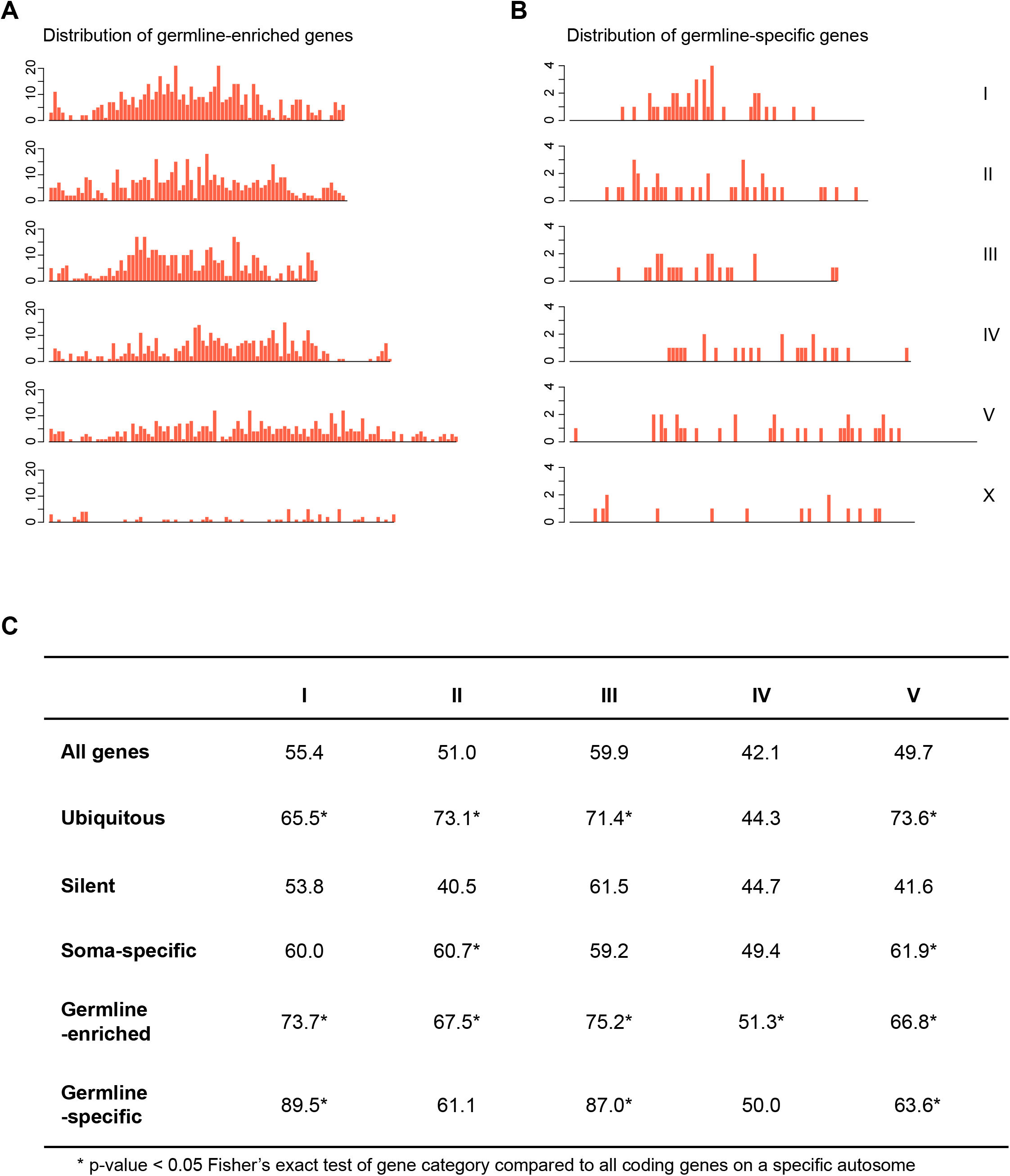
Chromosomal distribution of genes with germline expression. (A, B) Binned distribution of genes that show germline-enriched (A) and germline-specific (B) expression in 200 kb windows across the five autosomes and the X chromosome. (C) The percentage of genes in different categories of expression found in the centers of autosomes. Significant enrichment was determined by Fisher’s exact test between the number of genes of a given category in the center vs. arms of each autosome compared to all genes in the center vs. arms of the same autosome (*p-value < 0.05, Fisher’s Exact test).

**Figure S6:**
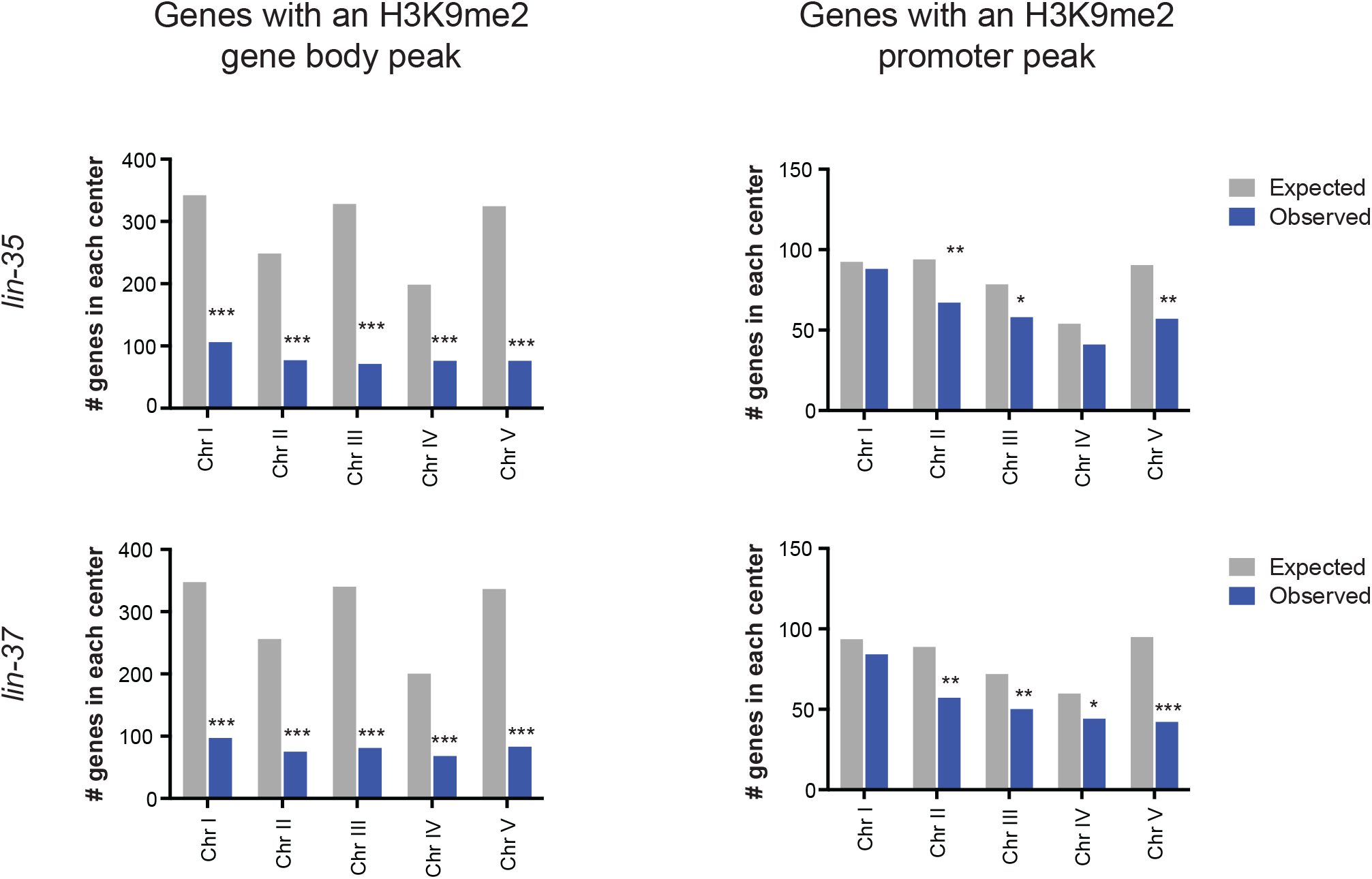
Chromosomal distribution of genes with H3K9me2 promoter or gene body peaks in *lin-35* and *lin-37* mutants. Enrichment analysis of genes with an H3K9me2 gene-body peak or promoter peak expected by chance and observed in chromosome centers *in lin-35* and *lin-37* mutant L1s. The expected number is based on the percentage of coding genes in the center versus arm regions of each chromosome; the observed number is the number of genes in the chromosome centers at 20°C. The locations of chromosome arm and center boundaries are from (Liu *et al.* 2010). Significant under-enrichment (black) was determined by the hypergeometric distribution (*p-value < 0.01, **p-value < 1 × 10^−5^, ***p-value < 1 × 10^−10^).

**Figure S7:**
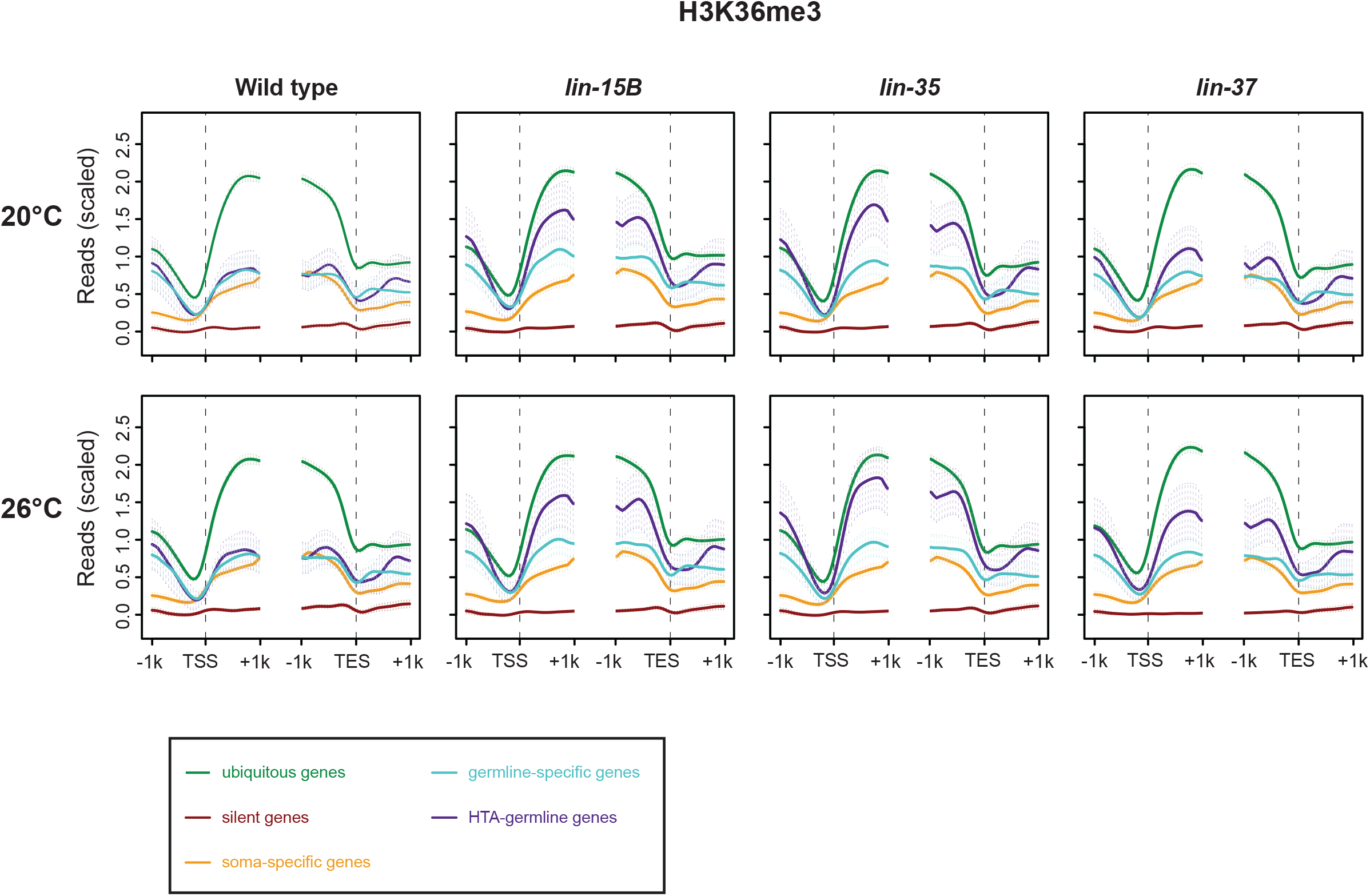
H3K36me3 over genes in wild-type and synMuv B mutant L1s. Metagene profiles of mean H3K36me3 ChIP-seq signal 1kb up- and downstream from the transcription start site (TSS) and transcription end site (TES) for genes that show ubiquitous, silent, soma-specific, germline-specific, or HTA-germline expression. Reads were scaled by dividing by the standard deviation and subtracting the 25th percentile. Error bars indicate 95% confidence intervals for the mean.

**Figure S8:**
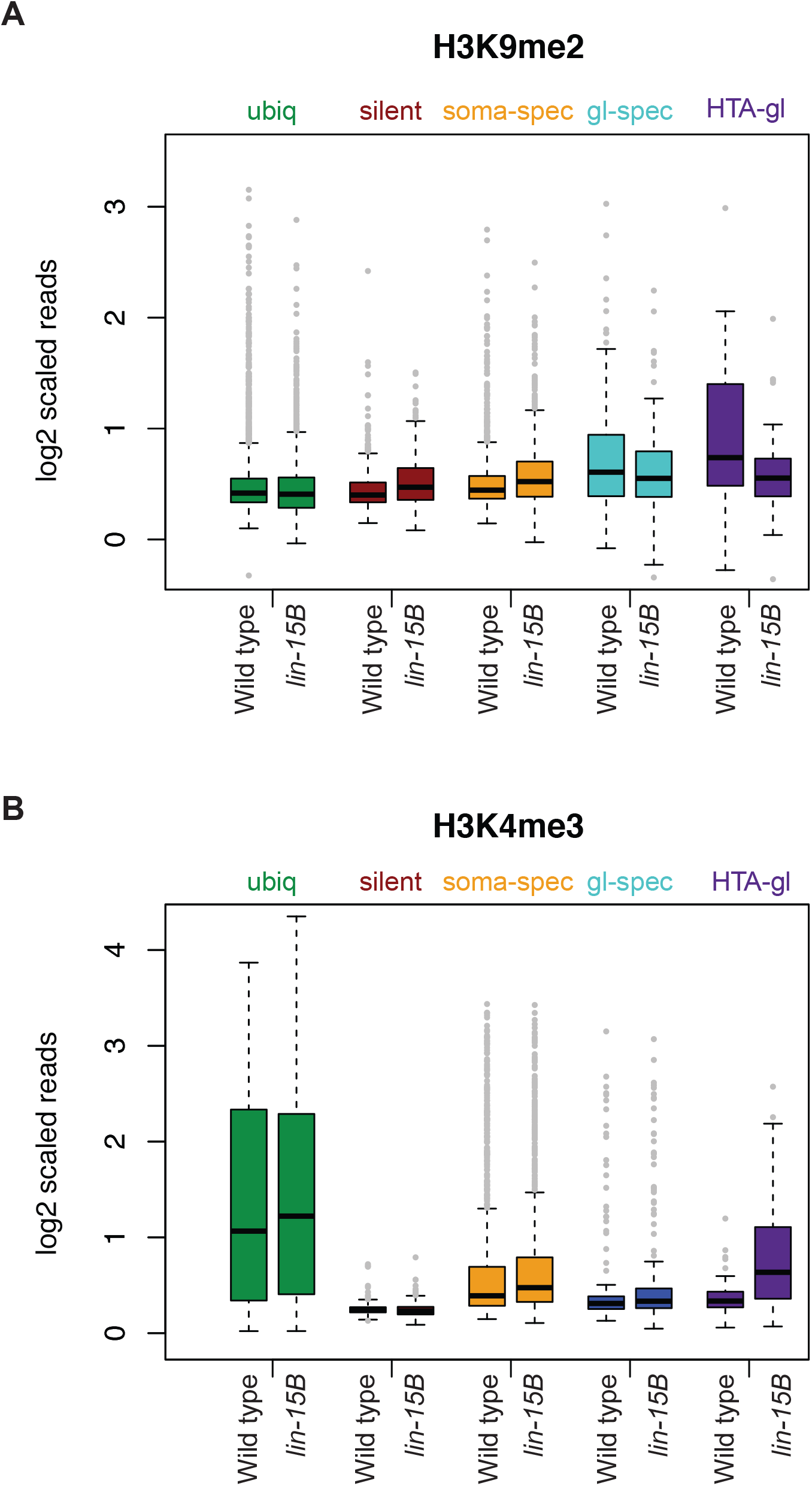
Comparison of H3K9me2 and H3K4me3 signal between wild-type and *lin-15B* mutant L1s. (A,B) Box-plots of H3K9me2 (A) and H3K4me3 (B) signal 250 bp up and downstream from the TSS in wild type and *lin-15B* mutants at 20°C for genes that show ubiquitous silent soma-spec¡f¡c germline_-_specific or HTA_-_germline expression

**Figure S9:**
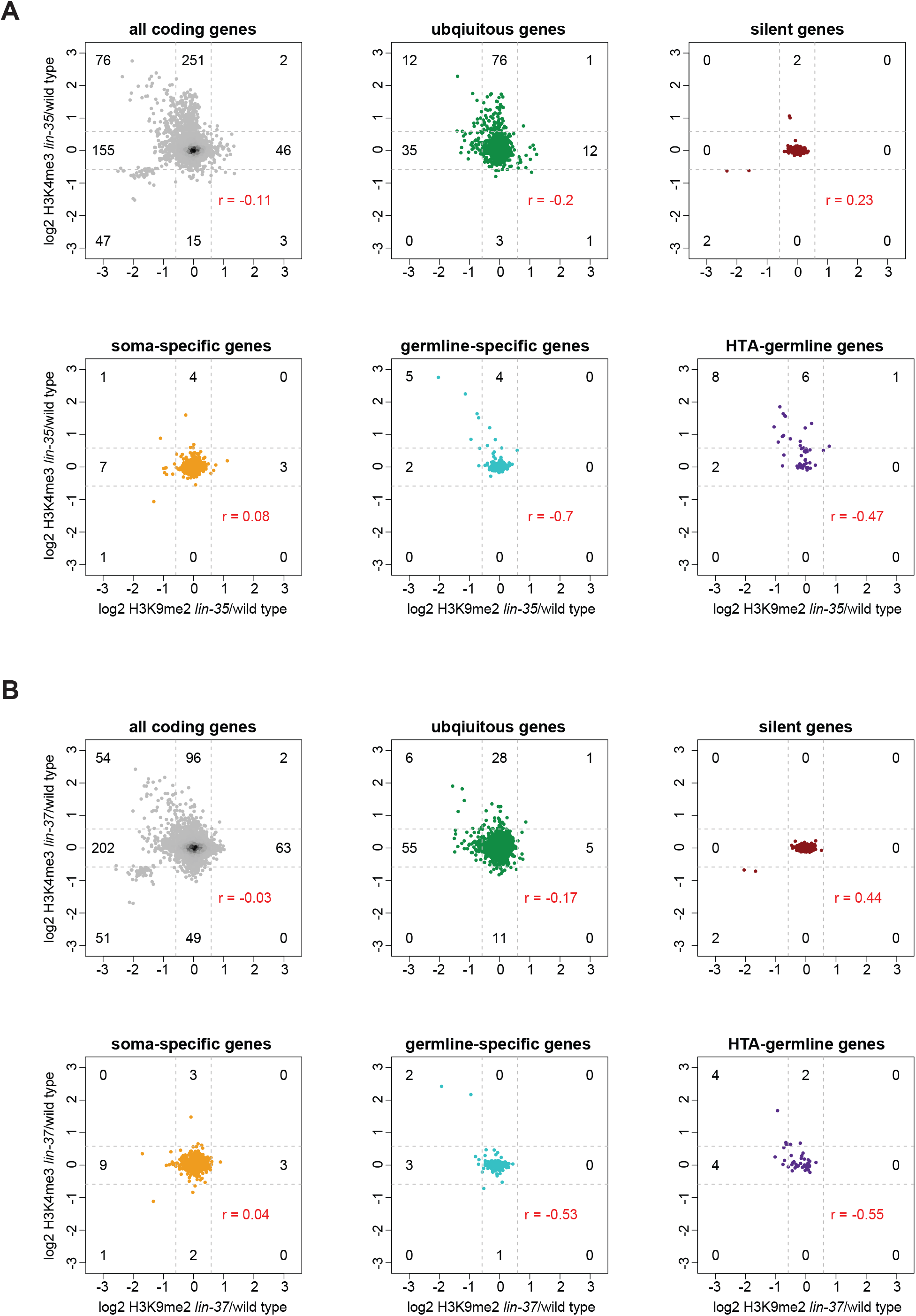
H3K4me3 vs H3K9me2 in *lin-35* and *lin-37* mutants. (A) Scatter plots of log2 fold change of the H3K9me2 signal in *lin-35* mutant/wild type vs. log2 fold change of the H3K4me3 signal in *lin-35* mutant/wild type within 250 bp upstream and downstream of the transcription start site (TSS) at 20°C for all coding genes and genes with ubiquitous, silent, germline-specific, soma-specific, or HTA-germline expression. Dotted lines represent 1.5 fold cut-offs; the numbers of genes above, below and within the cut-offs are indicated. r values show the Pearson correlation between changes in H3K9me2 and changes in H3K4me3 for each set of genes. (B) Similar analysis *for lin-37* compared to wild type.

